# How do avian embryos resume development following diapause? A new role for TGF-β in regulating pluripotency-related genes

**DOI:** 10.1101/2021.11.17.467607

**Authors:** Narayan Pokhrel, Olga Genin, Dalit Sela-Donenfeld, Yuval Cinnamon

## Abstract

Avian embryos can halt their development for long periods at low temperature in a process called diapause and successfully resume development when reincubated at maternal body temperature. Successful resumption of development depends on different factors, including temperature. We have recently shown that embryos that enter diapause at 18 °C present a significant reduction in their ability to develop normally when put back into incubation, compared to embryos entering diapause at 12 °C. However, the mechanisms underlying these differences are unknown. To address this question, transcriptome analysis was performed to compare the effect of diapause temperature on gene expression, and to identify pathways involved in the process. Genetic comparison and pathway-enrichment analysis revealed that TGF-β and pluripotency-related pathways are differentially regulated at the two temperatures, with higher expression at 12 °C compared to 18 °C. Investigating the involvement of the TGF-β pathway revealed an essential role for BMP4 in regulating the expression of the transcription factors Nanog and Id2, which are known to regulate pluripotency and self-renewal in embryonic stem cells. BMP4 gain- and loss-of-function experiments in embryos in diapause at the different temperatures revealed the main role of BMP4 in enabling resumption of normal development following diapause. Collectively, these findings identify molecular regulators that facilitate embryos’ ability to undergo diapause at different temperatures and resume a normal developmental program.

## Introduction

Fertilized avian embryos undergo several developmental transitions, including meroblastic cleavages, formation of the blastodisc, and compartmentalization into the inner disk—the area pellucida (AP) and the outer ring—the area opaca (AO) (Eyal-Giladi and Kochav 1976). At oviposition, embryos undergo blastulation, which is demarcated by the formation of epiblast and hypoblast cell layers. Based on the anterior expansion of the hypoblast, embryos are staged between stages X and XIII EG&K (Eyal-Giladi and Kochav 1976; Pokhrel et al. 2017). Remarkably, embryonic development at these stages can be suspended in response to low-temperature stimuli and the embryos enter into diapause. This dormancy period can last for weeks (Fasenko 2007). The ability to enter diapause has an essential evolutionary role in allowing for synchronized hatching of eggs, which are laid days apart in a single clutch. The diapause phenomenon is routinely exploited for commercial purposes to allow for egg storage prior to incubation (Brake et al. 1997).

Avian embryos are adapted to surviving during diapause and they can successfully resume development when placed back in incubation conditions. However, this phenomenon is highly dependent on a variety of factors, including environmental temperature and diapause duration (Pokhrel et al. 2018; Schulte-Drüggelte 2011). In particular, our recent finding demonstrated that embryos entering prolonged diapause (more than 7 days) at 12 °C or 18 °C retain much better cell viability, cytoarchitectural features and embryonic survivability at the lower temperature (Pokhrel et al. 2018; Pokhrel, Sela-Donenfeld, and Cinnamon 2021). Although the molecular mechanisms involved in this difference are unknown, it can be speculated that the lower temperature more efficiently reduces metabolic activity or activates specific signaling pathways, leading to better embryonic survivability following diapause.

Chicken blastodermal cells have been classified as embryonic stem cells (ESCs), and they share similar characteristics with mouse ESCs (Jean et al. 2015). Notably, in the latter, bone morphogenetic protein 4 (BMP4) has been found to play a crucial role in maintaining pluripotency and self-renewal capacity by promoting the expression of inhibitor-of-differentiation (Id) genes (Ying et al. 2003). *Id* genes are required to maintain cells in their stemness state (Hollnagel et al. 1999; Morikawa et al. 2016; Qi et al. 2004). In addition, the homeobox transcription factor Nanog has been identified as a key downstream effector of BMP4 signaling, again maintaining mouse ESC self-renewal and pluripotency (Kalmar et al. 2009; Pan and Thomson 2007; Torres and Watt 2008). Despite the known roles of BMP4, Id and Nanog in mammalian ESCs, it is not known whether they regulate the pluripotency state of the avian blastoderm, thereby promoting its ability to recover from diapause and resume normal development.

This study aimed to uncover the molecular mechanisms involved in the embryos’ ability to suspend their development for prolonged periods, and then successfully resume development. RNA-Seq of embryos which entered diapause at different temperatures and for different lengths of time revealed pathways and single genes involved in the divergence of their developmental trajectories when exiting diapause. We discovered that embryos placed in diapause at 12 °C halt their development and are better able to resume development than those entering diapause at 18 °C by maintaining expression of BMP4, which induced the expression of the ESC regulatory genes *Id2* and *Nanog*. This study sheds new light on the molecular pathways that act during diapause and preserve the embryos’ ability to successfully resume normal development following prolonged diapause.

## Results

### Transcriptome profiling of embryos in diapause at different temperatures and for different lengths of time

Avian embryos can initiate diapause at low temperatures and extend this process for prolonged duration. Moreover, they have the remarkable ability to successfully resume their development when incubated at maternal body temperature (~38 °C) (Figure 1A). We examined the effect of temperature and duration of diapause on the embryo’s ability to resume normal development. In practice, eggs can be stored within a range of storage temperatures, from 12 °C to 18 °C (reviewed in Pokhrel, Sela-Donenfeld, and Cinnamon 2021). We therefore stored the eggs at 12 °C or 18 °C for different lengths of time, and then incubated them at 37.8 °C for 21.5 days and monitored their ability to resume development and survive until hatch. Successful resumption of development was compared to development in fresh eggs that did not enter diapause prior to incubation. After 7 days of diapause at 12 °C or 18 °C, ~85% of the embryos survived regardless of diapause temperature, compared to 80% survival of the fresh embryos. However, beyond 7 days of diapause, the ability to successfully resume development diverged according to the temperature during diapause; 18 °C for 28 days led to a drastic drop in survival rate to ~15%, whereas at 12 °C, the survival rate declined only mildly, to ~70% (Figure 1A). Notably, most of the embryos that did not survive died within the first 3 days of incubation (Figure 1-figure supplement 1), suggesting that the early processes of blastulation, gastrulation and neurulation are the most strongly affected. Based on these results, the ability to successfully resume development following diapause was divided into two phases: an initiation period where the ability to resume development is maintained or slightly improved independent of the temperature, and an extension period where the ability to resume development is greatly dependent on the temperature during diapause.

**Figure 1.**
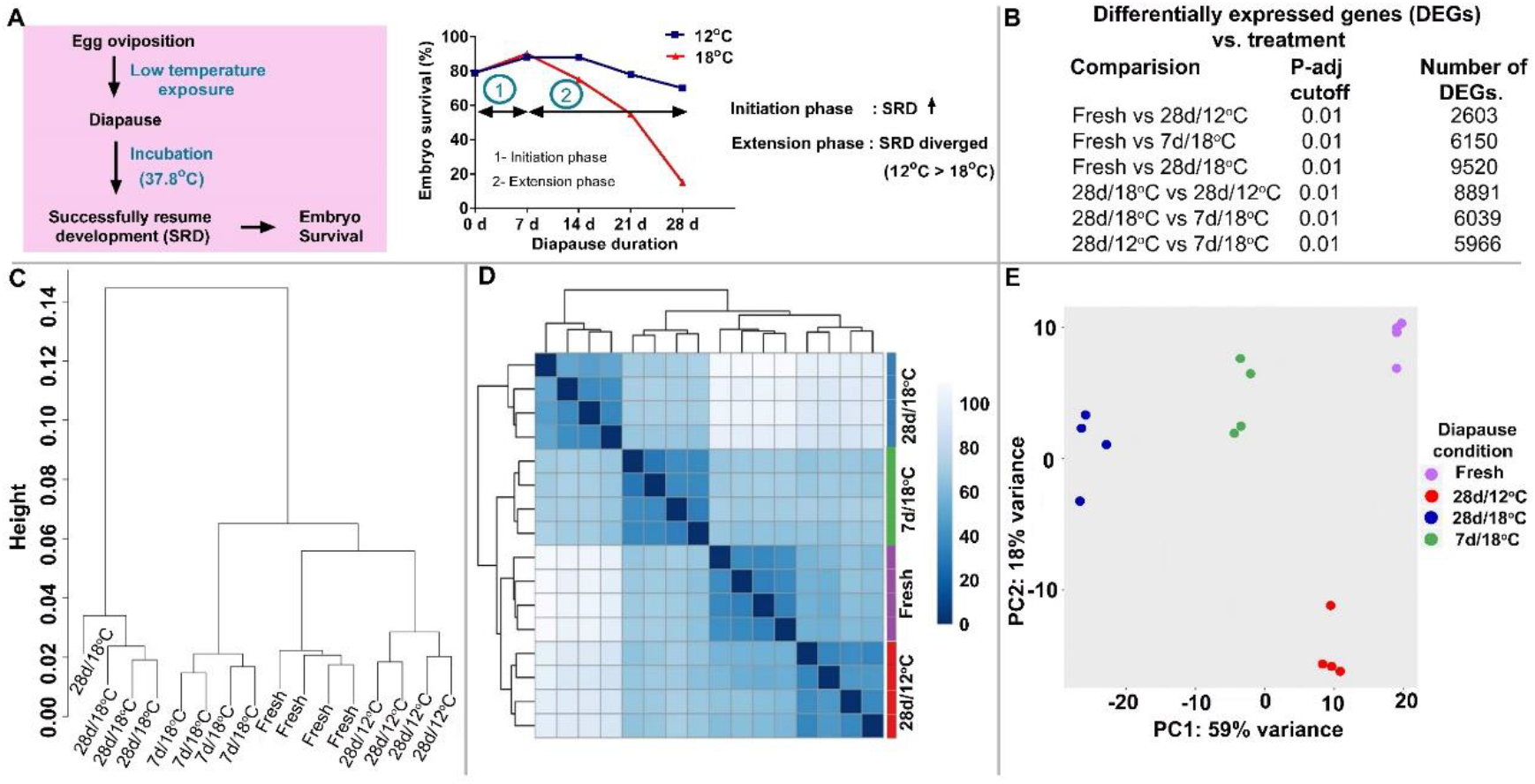
Embryo survival rate and gene-expression profile of embryos in diapause. (**A**) Embryos in diapause at low temperature for short or extended periods and then incubated at 37.8 °C show successful resumption of development (SRD), reflecting embryo survival; it increases during the initial 7 days of diapause (1) at both 12 °C and 18 °C, whereas beyond this point (extension phase of diapause, (2)), SRD is either maintained (at 12 °C) or declined (at 18 °C). (**B**) Differentially expressed genes (DEGs) in embryos under different diapause conditions (Fresh – no diapause; 28d/12°C – held in diapause for 28 days at 12 °C; 7d/18°C – for 7 days at 18 °C; 28d/18°C – for 28 days at 18 °C). (**C**) Hierarchical clustering of DEGs showing close clustering of fresh, 7d/18°C and 28d/12°C groups, compared to 28d/18°C group. (**D**) Correlation heat map. (**E**) PCA of the four embryo groups showing that the three groups fresh, 7d/18°C and 28d/12°C—are closely clustered, compared to the 28d/18°C group.

To decipher the molecular pathways underlying this phenomenon, RNA-Seq analysis was conducted under four different diapause conditions: *(i)* reference/control group of freshly laid eggs not subjected to diapause, *(ii)* eggs in diapause for 7 days at 18 °C (7d/18°C), *(iii)* eggs in diapause for 28 days at 12 °C (28d/12°C), and *(iv)* eggs in diapause for 28 days at 18 °C (28d/18°C). RNA-Seq analysis showed 6150 differentially expressed genes (DEGs) for the 7d/18°C group, 2603 DEGs for the 28d/12°C group, and 9520 DEGs for the 28d/18°C group, compared to the reference group (Figure 1B). Hierarchical clustering, correlation heat map, and principal component analysis (PCA) showed that the collection of genes expressed in the 28d/18°C group was most different from the fresh, 7d/18°C and 28d/12°C groups, whereas the latter three groups were correlated and closely clustered (Figure 1C-E). These data correlated with the embryonic survival phenotypes, where lower or higher number of DEGs between the groups was coupled with a maintained or reduced ability to successfully resume development, respectively.

### Different gene-regulatory networks are demonstrated in embryos in the extension phase of diapause at 12 °C vs. 18 °C

To identify pathways that might be involved in embryonic survivability during the extension phase, transcript profiles were compared between the 28d/12°C and 28d/18°C groups (Figure 2A). DEGs from these groups were analyzed for pathway enrichment using Gene Ontology (GO) molecular terms and Kyoto Encyclopedia of Genes and Genomes (KEGG) (Kanehisa 2019; Yi et al. 1999). This analysis revealed enrichment of pluripotency-related pathways, the TGF-β signaling pathway, and cancer-related pathway in the 28d/12°C group, and of apoptosis-related pathways, the Hedgehog signaling pathway, and proteasome pathway in the 28d/18°C group (Figure 2B). Specifically, *BMP4*, inhibitor-of-differentiation 2 (*Id2*), *Nanog, cPouV, Myc* and *Dazl* were upregulated in the 28d/12°C group, whereas *Gli1-3, Fas*, and *Pax6* were upregulated in the 28d/18°C group (Figure 2A).

**Figure 2.**
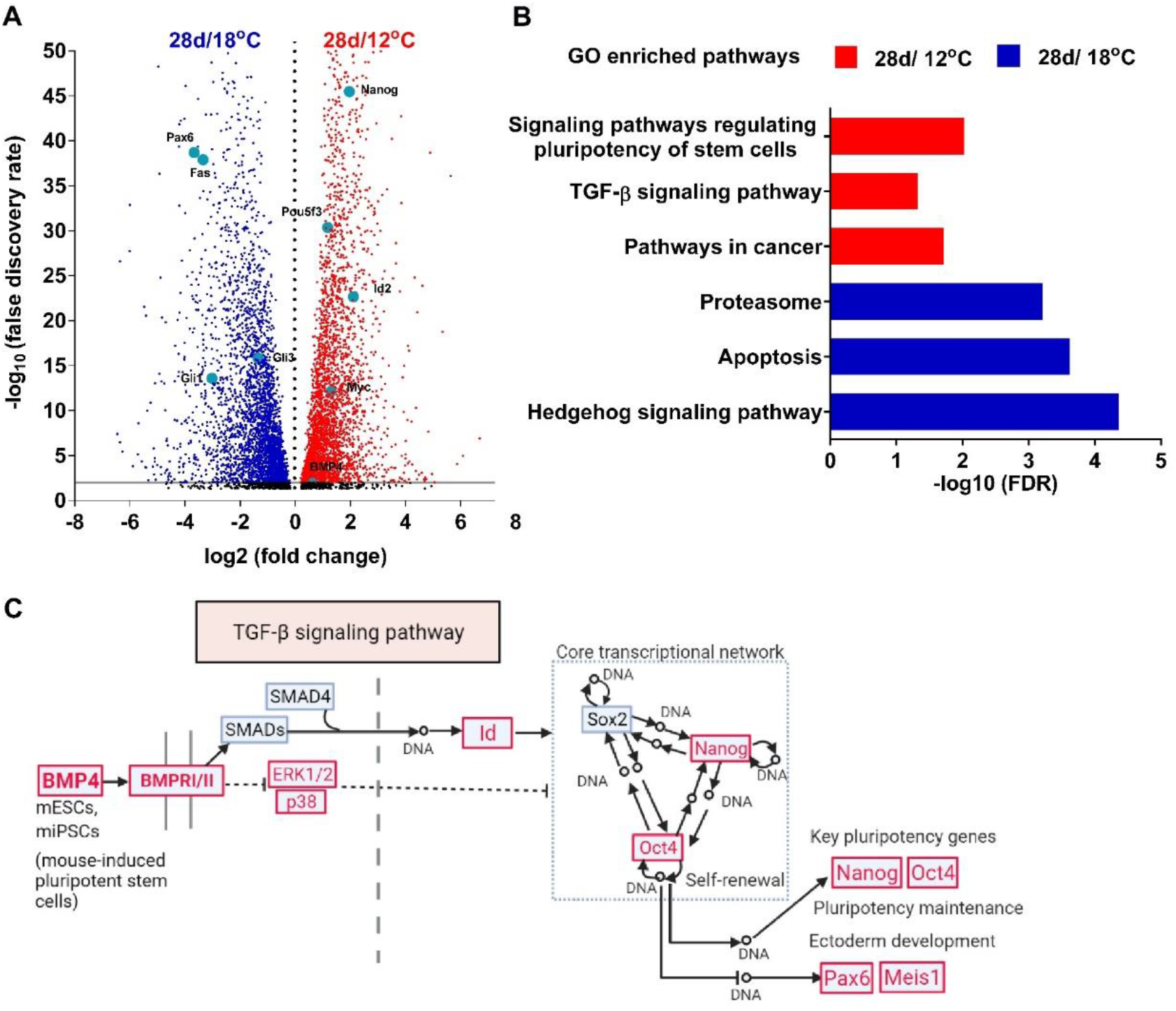
Gene-expression profile of embryos during the extension phase of diapause. (**A**) Volcano plot. Comparison of gene-expression profiles in 28d/18°C and 28d/12°C embryos. Upregulated genes in blue are from 28d/18°C group, and in red, from 28d/12°C group. (**B**) GO pathway-enrichment analysis of 28d/18°C and 28d/12°C groups using WebGestalt. Red and blue bars indicate enriched pathways in 28d/12°C and 28d/18°C groups, respectively. (**C**) Schematized road map for pluripotency regulation by the TGF-β pathway, based on KEGG pathway-enrichment analysis of the RNA-Seq data. Enriched gene sets are in red. An expanded scheme of enriched signaling pathways regulating pluripotency is shown in Figure 2-figure supplement 2.

### Downregulation of *BMP4* and pluripotency-related genes in embryos during the extension phase of diapause at 18 °C

In agreement with the enrichment of BMP4 and the pluripotency-associated genes *Nanog* and *Oct4* in the pathway analysis of the 28d/12°C group (Figure 2C), BMP4 was recently shown to positively regulate pluripotency genes in mouse ESCs (Ying et al. 2003; Yu et al. 2020). Therefore, we checked whether *BMP4* expression differs in embryos in diapause at 12 °C vs. 18 °C, and whether it induces the expression of pluripotency genes, which are essential for the ability to resume development. Further analysis of the RNA-Seq data showed that *BMP4* expression levels were already reduced after 7 days at 18 °C compared to the other groups, and were further downregulated after 28 days (Figure 3A). Concomitantly, expression of *Nanog* and *Id2*, previously shown to be downstream targets of *BMP4* (Luo et al. 2012; Ma et al. 2017; Xia et al. 2017), was also downregulated after 7 days of diapause at 18 °C (Figure 3A). To validate the RNA-Seq findings, real-time RT-PCR and whole-mount RNA in-situ hybridization (WMISH) analyses were performed on freshly laid embryos and eggs in diapause for 7 days or 28 days at 18 °C (Figure 3B-H). The quantified mRNA levels of both *Id2* and *Nanog* were ~3.5-fold lower in the 28d/18°C group compared to the control and 7d/18°C groups (Figure 3B). WMISH analysis further confirmed the broad expression of *Id2* and *Nanog* transcripts in the AO and AP of freshly laid embryos and 7d/18°C embryos (Figure 3C, D, F and G), while expression of both genes was markedly reduced in the blastoderm of the 28d/18°C group (Figure 3E and H). Notably, the expression of *Nanog*, but not *Id2*, was already slightly decreased after 7 days of diapause, indicating an earlier change in the former’s expression during the initiation phase of diapause. Together, the decreased expression of *BMP4, Id2* and *Nanog* during prolonged diapause at 18 °C suggested that the reduced embryonic survival following these diapause conditions is associated with a decrease in the pluripotency state of the blastoderm during prolonged diapause at relatively high temperatures.

**Figure 3.**
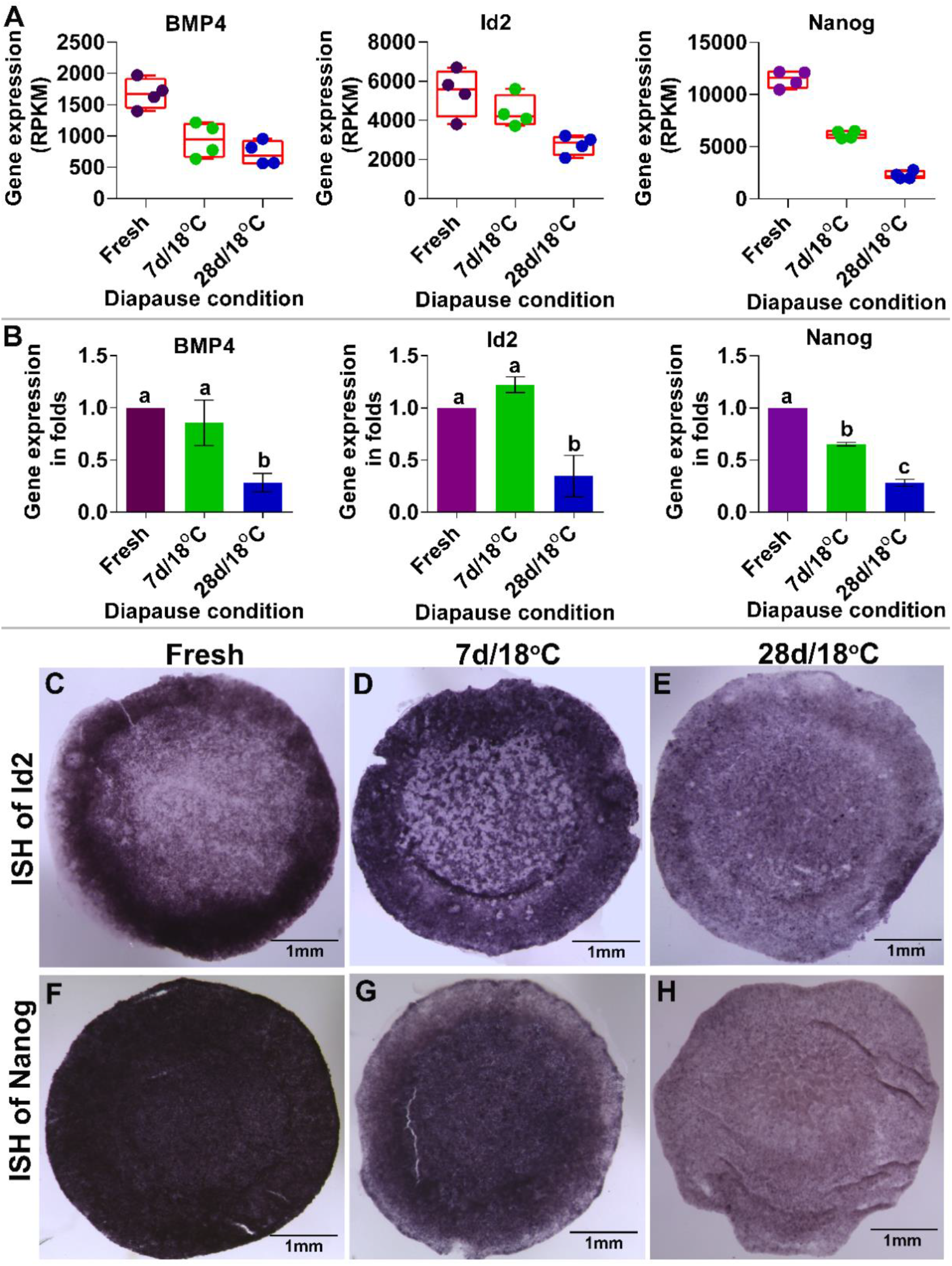
Analysis of BMP4-pathway activation under different periods of diapause at 18 °C. (**A**) Gene-expression profiles of *BMP4, Id2* and *Nanog* by RNA-Seq. (**B**) Quantitative validation of gene-expression analysis of *BMP4, Id2* and *Nanog* using real-time RT-PCR. Different letters above bars indicate significant difference between groups by ANOVA (*P* < 0.05). (**C-E**) WMISH of BMP4 target gene *Id2* in Fresh (**C**, n = 6), 7d/18°C (**D**, n = 6), 28d/18°C (**E**, n = 6) embryos. (**F-H**) WMISH of *Nanog* in Fresh (**F**, n = 6), 7d/18°C (**G**, n = 6), and 28d/18°C (**H**, n = 6) embryos.

### *BMP4* acts upstream of *Nanog* and *Id2*, and is sufficient to rescue their expression during diapause at 18 °C

Since *BMP4* expression was significantly reduced in embryos stored during the extension phase at 18 °C (Figure 3A and B), we next investigated whether the reduced expression of *Id2* and *Nanog* in the 28d/18°C group is a result of the reduced BMP levels, and can therefore be rescued by exogenous administration of BMP4. Blastoderms stored at 18 °C for 27 days were supplied with exogenous BMP4 for 24 h using two approaches: widespread application of recombinant BMP4 (200 ng/mL) to the entire blastoderm using pluronic gel (Makarenkova and Patel 1999), and local application using heparin beads soaked with BMP4 (Weisinger, Wilkinson, and Sela-Donenfeld 2008). Phosphate buffered saline (PBS) was added to the pluronic acid or beads as a control. First, to validate the beads’ BMP4-releasing efficiency, BMP4-soaked beads were applied in a separate experiment to the neural tube of 48-h-old embryos. This is known to accelerate neural crest delamination (Sela-donenfeld and Kalcheim 1999; Sela-Donenfeld and Kalcheim 2000), and was confirmed using HNK-1 immunostaining (Figure 4-figure supplement 3). Next, control or BMP4-treated blastoderms were collected and analyzed for *Nanog* and *Id2* mRNA expression using real-time PCR and WMISH. Real-time RT-PCR analysis of 28d/18°C embryos showed upregulation of *Id2* and *Nanog* transcript levels upon BMP4 application (Figure 4A) compared to controls. Notably, exogenous BMP4 was also found to induce its own expression in the 28d/18°C group (Figure 4A), suggesting that BMP4 levels are maintained in the embryos by a positive-feedback loop (Teo et al. 2012). These results were confirmed by WMISH, which showed upregulation of *Id2* and *Nanog* expression upon addition of BMP4 (Figure 4E-G) as compared to the PBS-treated embryos (Figure 4B-D). Specifically, whereas broad application of BMP4 using pluronic gel led to general upregulation of these genes in the AP (Figure 4E and F, respectively), implantation of BMP4-soaked beads resulted in their localized induction near the beads (Figure 4G, also see Figure 4D as control). Taken together, these results showed that BMP4 regulates the expression of the pluripotency-associated genes *Nanog* and *Id2* during diapause and is sufficient to restore the diminished expression levels seen during the extension phase of diapause at high temperature.

**Figure 4.**
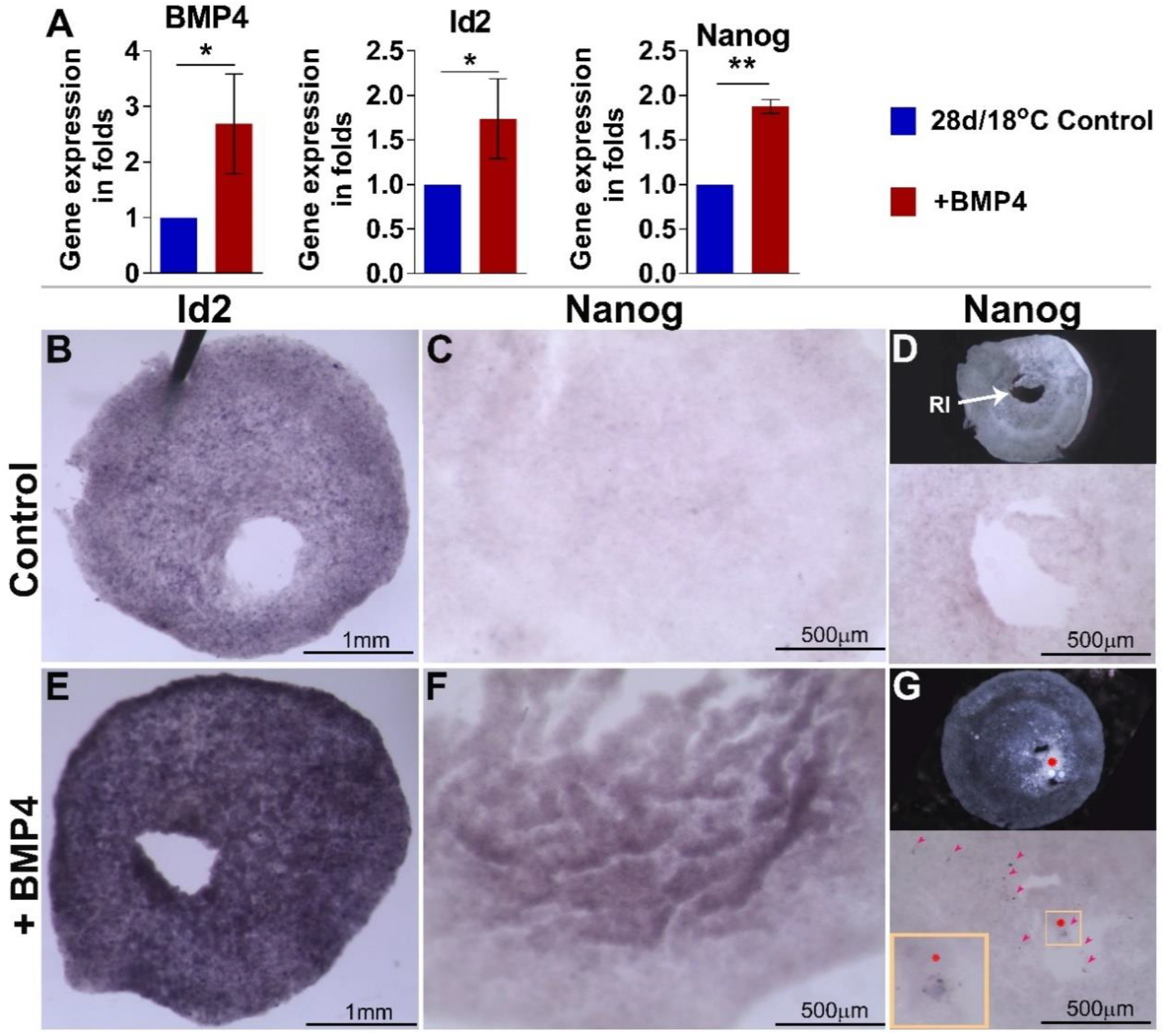
Ectopic application of BMP4 at the end of the extension phase of diapause at 18 °C induces the expression of pluripotency regulators *Nanog* and *Id2*. Embryos subjected to diapause for up to 27 days at 18 °C were treated with exogenous BMP4 recombinant protein for 24 h either globally using pluronic gel (**B, C, E, F**) or locally with beads (**D** and **G**). Following treatment, *Id2* and *Nanog* expression was analyzed by real-time RT-PCR and by probing their RNA expression using WMISH. (**A**) Exogenous treatment of BMP4 upregulates expression of *BMP4, Id2* and *Nanog* in 28d/18°C embryos. \**P* < 0.05; \*\**P* < 0.005; t-test. (**B** and **E**) WMISH of *Id2* gene expression in control (**B**, n = 4) and BMP4-treated (**E**, n = 4) 28d/18°C embryos; treatment was by pluronic gel. (**C** and **F**) WMISH of *Nanog* gene expression in control PBS-treated (**C**, n = 4) and BMP4-treated (**F**, n = 4) 28d/18°C embryos; treatment was by pluronic gel. WMISH of *Nanog* shows upregulated expression following BMP4 treatment (**F**). (**D** and **G**) WMISH of *Nanog* in PBS-treated control (**D**, n = 4) and BMP4-treated (**G**, n = 4) embryos; treatment was by heparin beads. Red asterisk denotes colocalization of *Nanog* expression with beads containing BMP4 protein. Red arrowhead represents BMP4-specific *Nanog*-expressing region. Inset in panel **G** (yellow square) shows induced *Nanog* expression near the transplanted BMP4-soaked beads at higher magnification. RI – hole in embryo showing region of injection for bead transplantation.

### Expression of *BMP4, Id2* and *Nanog* is maintained during the extension phase of diapause at low temperature

Our data revealed that prolonged diapause at 12 °C does not impair embryonic survival much when development resumes, in contrast to 18 °C (Figure 1A). This phenomenon correlates with the shared transcriptome profile of freshly laid eggs and 28d/12°C eggs, which differs from that of the 28d/18°C eggs (Figure 1C-E). Specifically, the RNA-Seq data showed largely similar expression of *BMP4* and *Nanog* and enhanced expression of *Id2* in the 28d/12°C group compared to the reference group (Figure 5A), in contrast to their profound reduction in the 28d/18°C group (Figure 3A). To further validate these results, real-time RT-PCR was performed on blastoderms collected from freshly laid eggs and from those of the 28d/12°C group. Quantification of the mRNA levels of these genes confirmed the RNA-Seq results, showing that the expression of *BMP4* and *Nanog* does not differ significantly between the 28d/12°C and reference groups, whereas *Id2* levels increased in the latter embryos (Figure 5B). WMISH analysis was also performed on embryos stored at 12 °C for 7 or 28 days, revealing no decrease in the expression of *Id2* and *Nanog* during the extension phase (Figure 5C-F), in contrast to their decline in the 28d/18°C group (Figure 3C-H). These results indicated that expression of *BMP4, Nanog* and *Id2* is maintained in embryos subjected to diapause at 12 °C during both initiation and extension phases, as opposed to their downregulation in embryos in diapause at 18 °C for up to 28 days.

**Figure 5.**
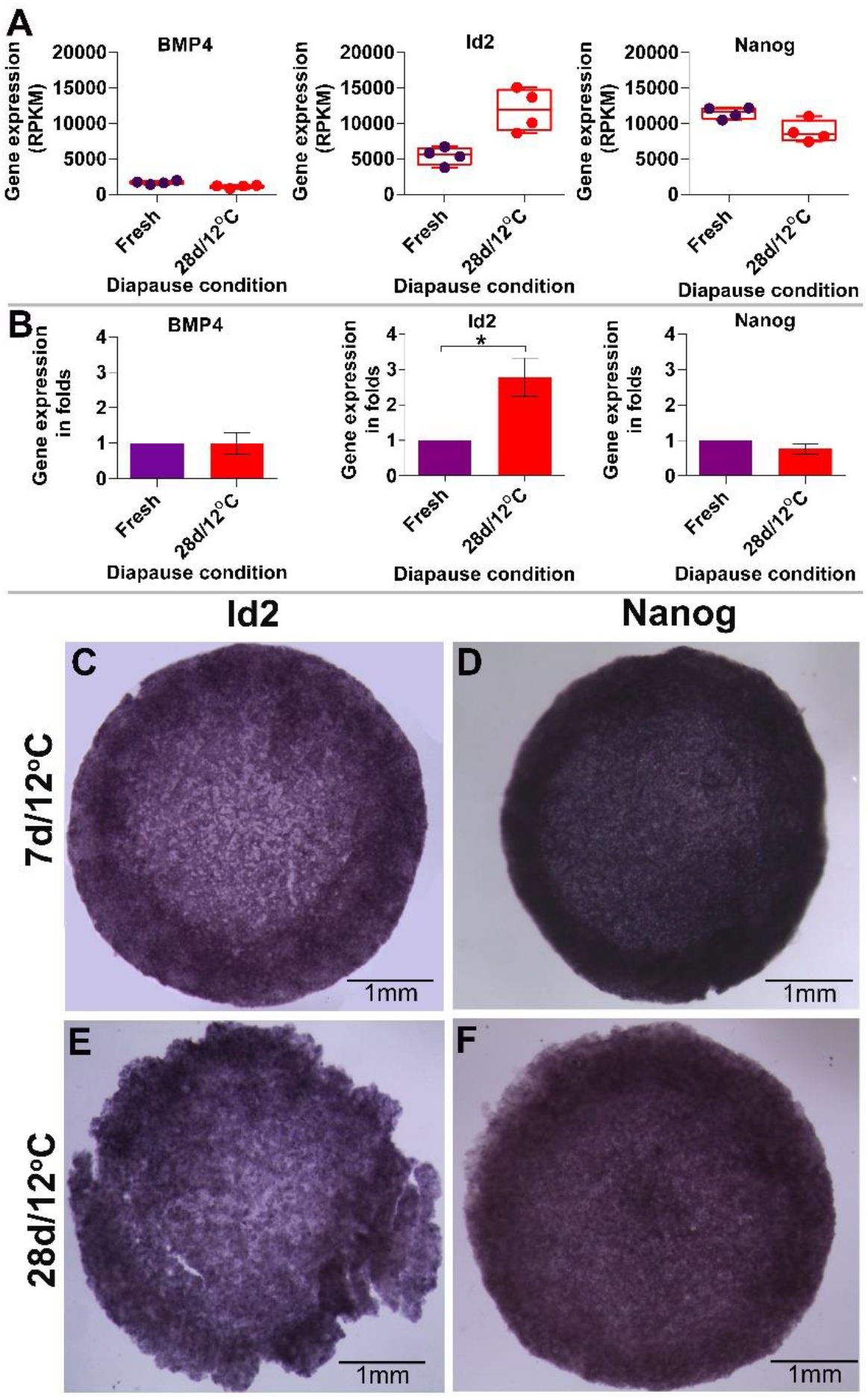
Embryos in diapause at 12 °C maintain expression of *BMP4, Id2* and embryonic pluripotency regulator *Nanog*, throughout the extension phase. (**A**) RNA-Seq profile of *BMP4, Id2* and *Nanog* expression in fresh and 28d/12°C embryos. These three genes were maintained in embryos throughout the extension phase of diapause at 12 °C. (**B**) Quantitative validation of *BMP4, Id2* and *Nanog* expression using real-time RT-PCR. \**P* < 0.05 by t-test. (**C** and **E**) WMISH analysis showing maintained expression of BMP4 target gene *Id2* in 7d/12°C (**C**, n = 6) and 28d/12°C (**E**, n = 6) embryos. (**D** and **F**) WMISH analysis showing maintained expression of *Nanog* in 7d/12°C (**D**, n = 6) and 28d/12°C (**F**, n = 6) embryos.

### Treatment with the BMP4 inhibitor Noggin results in downregulation of *Id2* and *Nanog* in embryos in diapause at 12 °C

The maintained expression of *BMP4, Id2* and *Nanog* during the extension phase at 12 °C, but not 18 °C, correlates with the embryos’ ability to resume development (Figure 1A, 3, 5). Since BMP4 was found to be upstream of *Id2* and *Nanog* and to restore their reduced expression in embryos that stayed in prolonged diapause at 18 °C (Figure 4), we next asked whether inhibition of BMP4 would result in their downregulation in the 7d/12°C group. BMP4 was inhibited by applying Noggin, an extracellular soluble BMP inhibitor (Di-Gregorio et al. 2007; Harland 2000; Joubin and Stern 1999; Weinstein and Hemmati-Brivanlou 1997) to the embryos in diapause. Freshly laid embryos were treated with conditioned media from control or Noggin-producing Chinese hamster ovary (CHO) cells (Figure 6-figure supplement 5) (Lamb et al. 1993; Sela-Donenfeld and Kalcheim 2002) mixed with pluronic gel, and stored for 7 days at 12 °C. WMISH analysis was performed to determine *Id2* and *Nanog* expression (Figure 6). Whereas control embryos expressed *Id2* and *Nanog* at levels similar to those in the untreated embryos (compare Figure 6A and C with Figure 5C and D, respectively), Noggin-treated embryos showed a marked reduction in both genes’ expression (Figure 6B and D), further confirming that BMP4 is upstream of these genes and suggesting that it is essential to maintaining their pluripotency potential during diapause.

**Figure 6.**
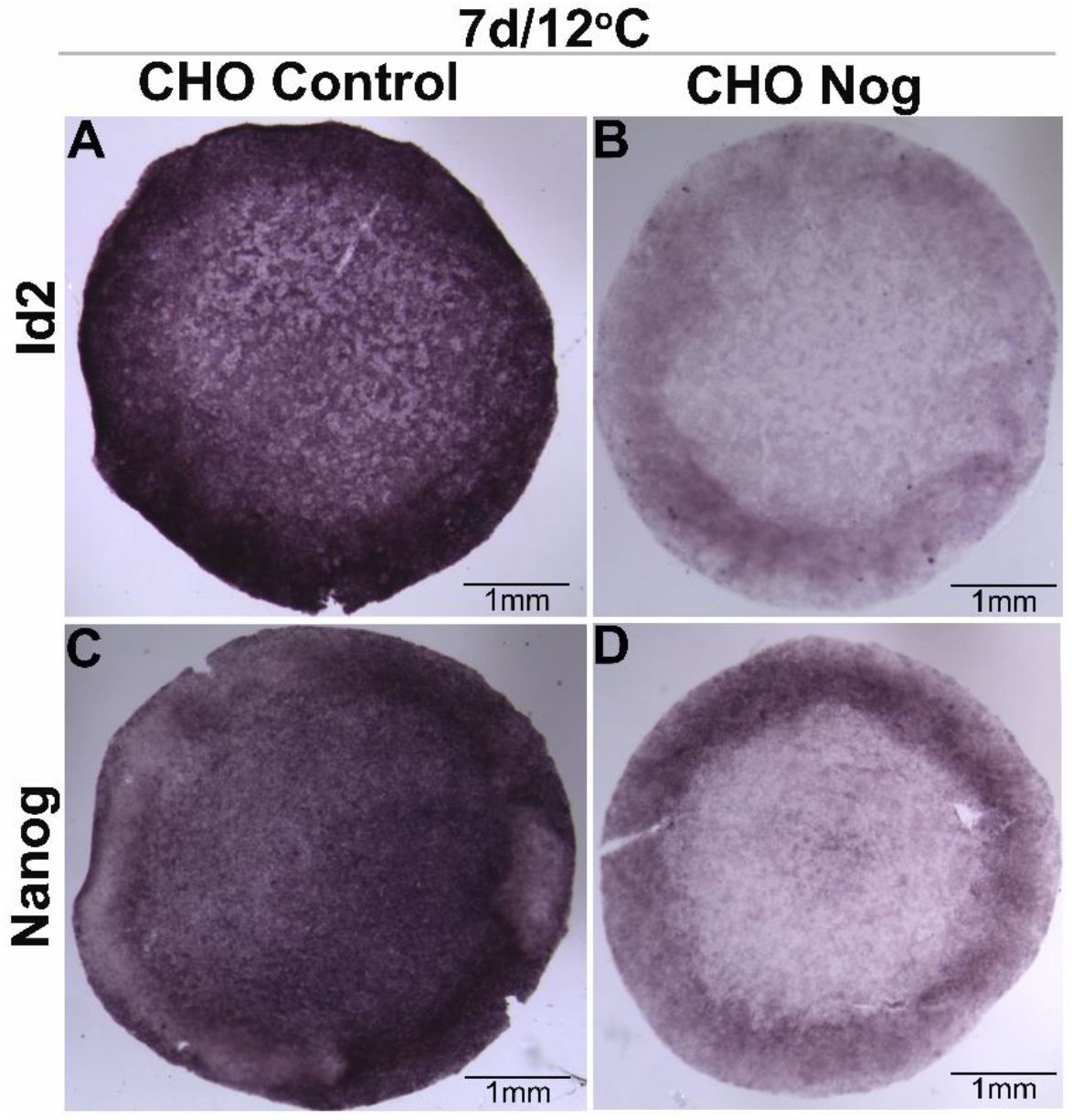
Treatment with the BMP4 inhibitor Noggin results in downregulation of *Id2* and *Nanog* in embryos in diapause at 12 °C. (**A** and **B**) Fresh embryos were treated with CHO-conditioned (CHO Control) or CHO-Noggin-conditioned (CHO Nog) media and then placed into diapause for 7 days at 12 °C and subjected to WMISH analysis of *Id2*. (**A**) CHO Control (n = 6). (**B**) CHO Nog (n = 6). CHO Nog-treated embryos showed decreased *Id2* expression, whereas in CHO Control-treated embryos, its expression remained unaffected. (**C** and **D**) WMISH of *Nanog* (n = 6) in treated embryos subjected to diapause for 7 days at 12 °C. (**C**) CHO Control. (**D**) CHO Nog. CHO Nog-treated embryos showed decreased *Nanog* expression, whereas in CHO Control-treated embryos, its expression remained unaffected.

### Expression of key pluripotency-related genes is maintained in embryos in diapause for a prolonged time at low temperature

To further determine whether the expression of pluripotency-related genes is better maintained at 12 °C than at 18 °C, the expression of other key pluripotency marker genes, *Dazl, cPouV*, and *Myc*, was analyzed by RNA-Seq. These genes were downregulated in the 28d/18°C group relative to the fresh blastoderms, and to the 28d/12°C and 7d/18°C groups (Figure 7A). This was supported by the PCA which showed clustering of the pluripotency-related genes in the fresh eggs, and 7d/18°C and 28d/12°C groups, while the 28d/18°C group was distantly clustered (Figure 7B). Notably, while embryos of the 28d/18°C group downregulated pluripotency markers, markers that are associated with more advanced developmental stages were upregulated, such as the neuronal cell fate marker *Pax6* (Figure 7C). This suggested that embryos in diapause at 18 °C for more than 7 days are advancing toward a more differentiated state. To fully validate these results, *Pax6* expression levels were determined by real-time RT-PCR in all embryonic groups (Figure 7D). *Pax6* expression was significantly enhanced in the 28d/18°C group compared to the other groups. Taken together, these results suggested that the BMP4 signaling pathway, which induces the expression of pluripotency-related genes, is maintained during the initiation phase of diapause regardless of egg temperature, and is associated with the ability to resume development after diapause. However, during the extension phase at the higher temperature, there is a marked reduction in BMP4 and pluripotency genes that is coupled with premature onset of neural differentiation, which may be linked to the embryos’ poor survival following diapause.

**Figure 7.**
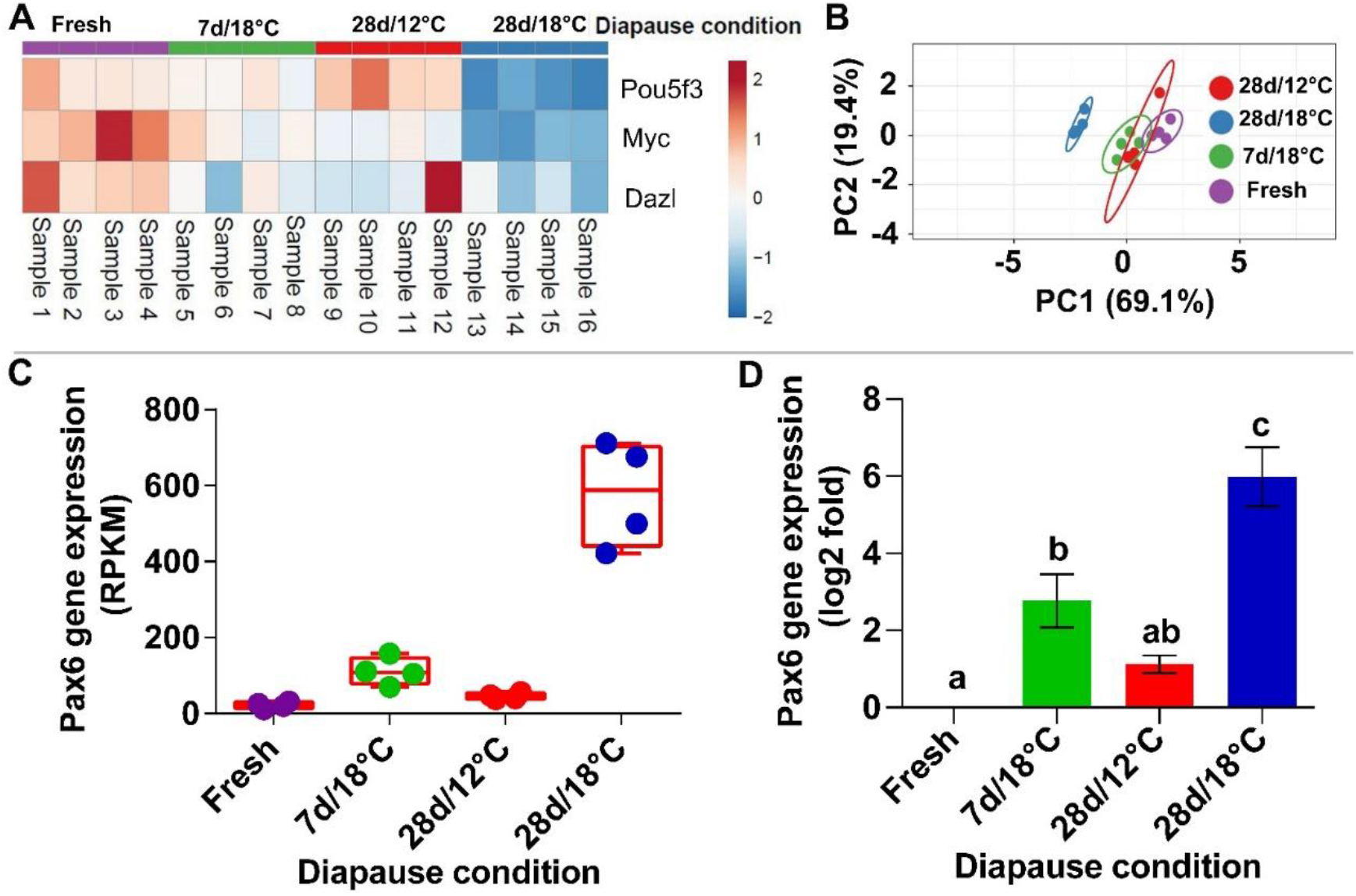
Expression profile of genes involved in pluripotency and differentiation in embryos during diapause. (**A**) Heat map of genes involved in pluripotency revealed a decrease in their expression in the 28d/18°C group, compared to the other three groups (fresh, 7d/18°C and 28d/12°C). (**B**) PCA of genes involved in pluripotency revealed close clustering of pluripotency-associated genes in fresh, 7d/18°C and 28d/12°C groups, whereas the 28d/18°C group was distantly clustered. (**C** and **D**) RNA-Seq expression profile of *Pax6* in embryos in diapause (**C**), and validation using real-time RT-PCR (**D**) showed upregulation in its expression following 28 days of diapause at 18 °C. Different letters above a bar indicate significant difference between groups by ANOVA (*P* < 0.05).

### BMP4 enhances successful resumption of embryo development after prolonged diapause at 18 °C

As the ability to resume development is significantly compromised following prolonged diapause at 18 °C and is coupled with decreased expression of BMP4 and pluripotency-related markers, we next sought to determine whether exogenous addition of BMP4 can enhance embryonic survival following diapause at 18 °C. Embryos that had been in diapause for 11 days at 18 °C were treated with PBS (mock) or BMP4-containing pluronic acid, and diapause was continued for 3 more days under the same conditions, followed by incubation for 72 h at 37.8 °C before assessing the embryos’ developmental state (Figure 8A). The rationale for testing embryos after 14 days in diapause (14d/18°C) was that this is the time point at which the decrease in the ability to resume development begins to be evident. As a positive control, an additional group of 7d/18°C embryos, which have a high survival rate (Figure 1A), was treated with PBS and directly incubated for 72 h at 37.8 °C (Figure 8B). All embryos were evaluated for their development (Figure 8C-E), and based on their morphology and survival rates, they were divided into three groups: *(i)* failure to successfully resume development, which was defined as fertile embryos which failed to form body axis and blood islands, *(ii)* a mild recovery phenotype with embryos that displayed body axis and formation of blood islands, but had no overt beating heart, and *(iii)* successful recovery, in which embryos developed brain vesicles, eyes, closed neural tube, beating heart with blood circulation and somites. These parameters were also quantified and summed as final scores (Figure 8F), based on the different criteria (Figure 8-figure supplement 4).

**Figure 8.**
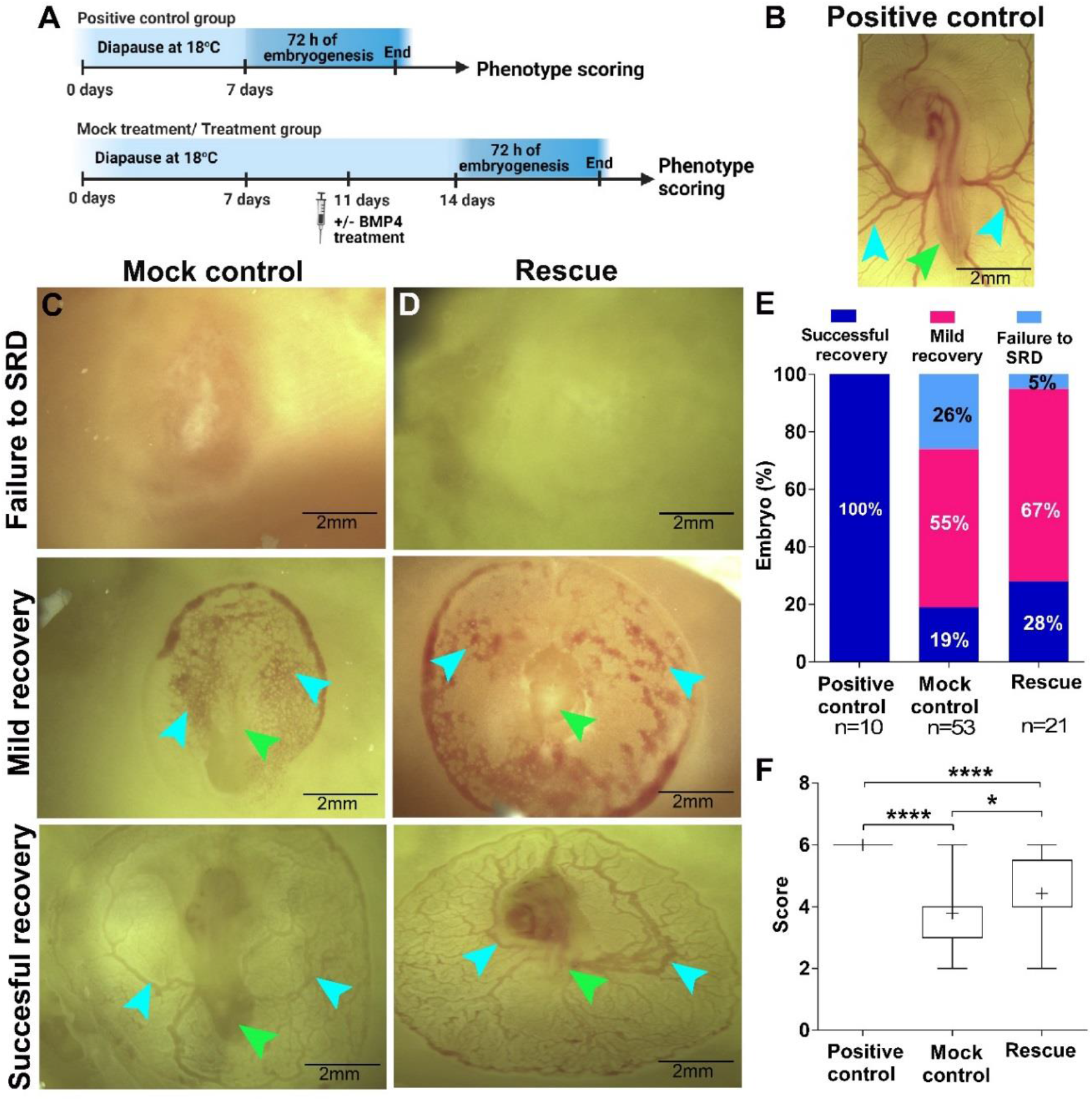
BMP4 enhances successful resumption of development (SRD) in embryos in prolonged diapause at 18 °C. (**A**) Schematic representation of experimental design. Embryos that underwent diapause up to the extension phase (11d/18°C) were treated with recombinant BMP4 protein for 3 days under the 18 °C diapause condition. Pluronic gel mix without BMP4 protein applied to embryos under the same diapause conditions served as a mock control. Embryos that underwent only the initiation phase of diapause (7d/18°C, without any BMP4 treatment) were used as positive controls as they have more SRD following this diapause condition. Embryos from all groups were incubated for 72 h and their phenotypes were scored. (**B**) All of the embryos from the positive control group showed SRD following diapause. The embryos had well-developed blood vessels (blue arrowhead) and embryo body axis (green arrowhead). (**C** and **D**) Phenotypes of positive control, mock control and rescued embryos were defined by three categories: (i) failure to SRD (no SRD characteristics), (ii) mild recovery, and (iii) SRD. Interestingly, embryos with the SRD phenotype from the BMP4-treated rescue group showed more branching of blood vessels (**D**, blue arrowhead) and smaller axis formation (**D**, green arrowhead), compared to positive controls (**B**) and mock controls (**C**). (**E**) Phenotype distribution of embryos. All positive control embryos showed the SRD phenotype. Compared to the mock control (**C**), a higher number of embryos had the SRD phenotype following BMP4 application (rescue, **D**) during the extension phase of diapause. (**F**) Phenotype score value (based on embryo morphology) of embryos following 72 h of embryogenesis. Each successive development event in post-diapause embryos (positive control, mock control and BMP4-treated rescue group) were scored from 1 to 6 (Figure 8-figure supplement 4) and the cumulative value was analyzed. A higher cumulative score represents better embryogenesis post-diapause. Wilkinson box-plot analysis shows higher score values in BMP4-treated rescue embryos than in mock controls, suggesting significant rescue of the SRD phenotype following treatment with BMP4. \**P* < 0.05, \*\*\*\**P* < 0.0001; t-test. Bars show the upper and lower limit of the score value.

Our results showed that 100% of the PBS-treated embryos from the 7d/18°C group displayed successful recovery (Figure 8B and E), whereas in the PBS-treated embryos of the 14d/18°C group, only 19% fully recovered, 26% failed to successfully resume development and 55% showed the mild recovery phenotype (Figure 8C and E). In contrast, in the BMP4-treated group, only 5% of the embryos failed to successfully resume development, 67% showed the mild recovery phenotype, and 28% recovered fully after 72 h of incubation (Figure 8D and E). In addition, scoring of the 14d/18°C embryos further confirmed the higher recovery of the BMP4-treated embryos compared to the PBS/mock controls (4.4 + 0.2 compared to 3.7 + 0.1, respectively, Figure 8F). Collectively, this experiment showed that addition of BMP4 is sufficient to partially rescue the loss of survival following diapause during the extension phase at 18 °C. Based on our findings, we propose a model in which BMP4 plays a central role in allowing embryos to successfully resume development following prolonged diapause at low temperature by maintaining the expression of pluripotency genes; this role is impaired during prolonged diapause at higher temperature (Figure 9).

**Figure 9.**
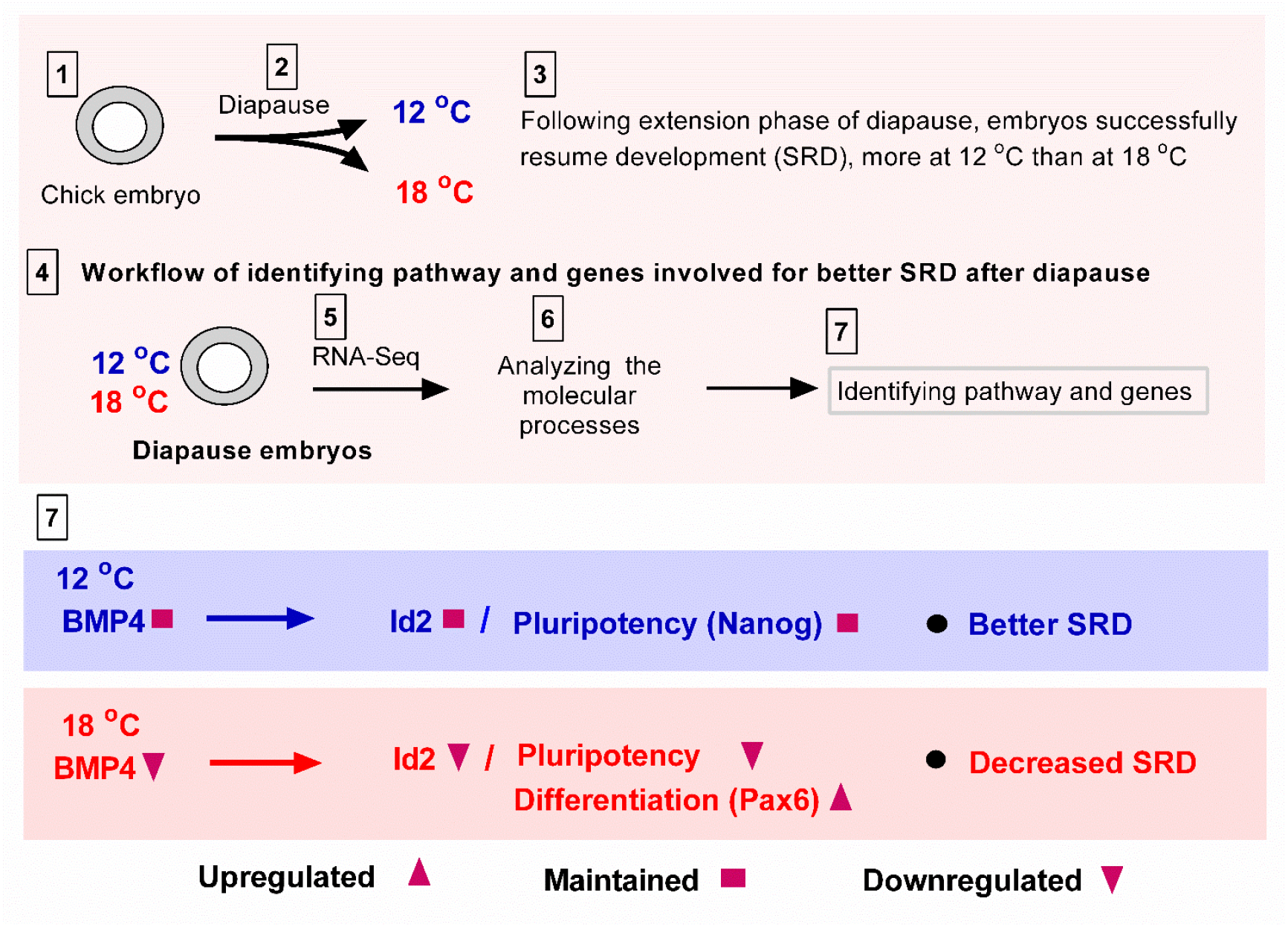
Summary of molecular mechanism involved in embryo survival during and after diapause. Embryos can enter diapause at low temperature and successfully resume development (SRD). SRD improves during the diapause initiation phase, independent of diapause temperature. However, beyond the initiation phase, in the extension phase, SRD diverges depending on diapause temperature: it is maintained at 12 °C and decreased at 18 °C. To understand this divergence, the pathway and genes were analyzed. RNA-Seq results revealed a role for BMP4 in the positive regulation of the pluripotency-related gene *Nanog* during diapause, and for lineage commitment after diapause.

## Discussion

### Initiation and extension phases of embryonic diapause

This study reveals the gene regulatory networks that participate in early and late stages of avian embryo diapause. Our recent finding of a correlation between diapause conditions and embryonic ability to successfully resume development (Pokhrel et al. 2018), together with our transcriptome analysis and validation of molecular processes in this study, suggest that diapause can be temporally subdivided into two phases: an initiation phase, where the ability to resume development is maintained or slightly improved at lower and higher temperatures, and an extension phase where the ability to resume development is temperature-dependent, largely decreasing with increasing temperature. This finding corroborates previous studies which showed that subjecting ovipositioned eggs to diapause for several days is beneficial for resumption of development, as compared to freshly laid eggs, and suggested that this results from the transition from hypoxic conditions prior to oviposition to normoxic conditions thereafter (Akhlaghi et al. 2013; Fasenko 2007; Karoui et al. 2006; Lapao, Gama, and Soares 1997). Moreover, the decline in the ability to resume development following longer diapause at higher temperatures also confirms previous studies which showed that each day of diapause beyond 7 days at 18 °C leads to a striking decrease in development resumption (Brake et al. 1997; Tullett 2009).

To determine the molecular processes responsible for these phenomena, RNA-Seq was performed on embryos in diapause for different lengths of time at different temperatures. Changes in gene-expression profiles were already seen in the initiation phase, although they did not lead to maladaptive changes in development resumption. Hence, these transcriptional changes are likely to result from the transition of embryos from a ‘non-diapause state’ prior to oviposition, to a diapause-induced state, at both 12 °C and 18 °C. In contrast, embryos in the extension phase of diapause demonstrated dramatic differences in gene-expression profiles according to the temperature, which was tightly linked with the ability to successfully resume development; the gene-expression profile was significantly different in embryos that underwent diapause for 28 days at 18 °C vs. 12°C, and was closely associated with poor ability of the former, but not the latter, to successfully resume development. PCA further showed that the gene-expression profiles of fresh embryos and those in diapause for 7 days at 18 °C and for 28 days at 12 °C are closely clustered, whereas genes expressed in embryos in diapause for 28 days at 18 °C were distantly clustered from all other groups. These findings strongly suggest that embryos in diapause for a long period of time at low temperature adapt better to resumption of development due to their ability to maintain the expression of central genes, an ability which is absent at higher temperatures. Our findings also highlight the marked plasticity of blastodermal cells to recover from undesirable conditions, which is likely to provide an evolutionary advantage for bird and reptile embryos which are much more exposed to environmental changes than mammalian embryos. Another example of this phenomenon is the utilization of a protocol in which stored eggs are put into a short period of incubation and then put back into storage, reducing the detrimental effect of egg diapause (Dymond et al. 2013) although in that study, the molecular mechanisms underlying the reactivation of biological processes by this type of storage modification were not known.

### Roles of the BMP4 pathway in maintaining embryos’ ability to successfully resume development

Which molecular mechanisms participate in maintaining embryos’ ability to resume development following the extension phase of diapause at 12 °C, but not at 18 °C? Our data revealed that BMP4 expression is maintained at 12 °C, compared to 18 °C, and that it positively regulates the pluripotency-related genes *Nanog* and *Id2*. BMP-dependent induction of these genes may be required as an adaptive mechanism for embryos in diapause to resume normal development following incubation. Previous studies in chicken embryos have shown that BMP4 is expressed in the blastoderm, and that during early gastrulation, its expression is downregulated in the AP but maintained in the AO (Streit et al. 1998). At later stages, the expression of several BMP4-signal inhibitors, such as Noggin and Chordin, is initiated in the anterior primitive streak and notochord, to allow neurulation (Di-Gregorio et al. 2007; Harland 2000; Streit et al. 1998; Weinstein and Hemmati-Brivanlou 1997; Wittler and Kessel 2004; Zimmerman, De Jesús-Escobar, and Harland 1996). These findings indicate that BMP4 signaling is activated prior to gastrulation and has to be blocked to allow further development of the gastrulating embryo. Notably, chicken blastoderm cells have been suggested to be equivalent to mouse ESCs (Jean et al. 2015). Since several pieces of evidence have indicated that BMP4 signaling is active in mammalian ESCs, maintaining their pluripotency (Morikawa et al. 2016; Qi et al. 2004), we suggest that BMP4 signaling plays a similar role during diapause at low temperature to maintain the pluripotency state of blastoderm cells, whereas at higher temperatures, this signaling is less active, leading the cells to wrongly proceed in their development, as reflected by the upregulation of the neural development gene *Pax6*. Direct support for this possibility was provided by the ability to partially rescue the development of embryos in diapause at 18 °C by application of BMP4, which led to the induction of *Nanog* and *Id2* expression and to the ability to develop beyond gastrulation and give rise to a normal or less perturbed phenotype in comparison to untreated embryos. However, further study is required to unravel the molecular machinery downstream of BMP4 that induces the expression of the *Id2* gene, as other pathways, including Fgf and Egf signaling, are also known to promote *Id* gene expression in other contexts (Lasorella, Benezra, and Iavarone 2014; Roschger and Cabrele 2017).

In-silico pathway analysis of the embryos that underwent diapause at 18 °C for an extended period revealed that downregulation of BMP4 and pluripotency-related gene targets was coupled with enrichment of the Hedgehog signaling pathway and premature expression of Pax6 which, under normal conditions, are expressed in more advanced embryos (Bertrand, Medevielle, and Pituello 2000; Teixeira et al. 2018). In addition, enrichment of the apoptosis pathway was demonstrated in this group, in agreement with previous studies that demonstrated increased cell death during diapause at 18 °C (Bakst and Akuffo 2002; Pokhrel et al. 2018). A central remaining question is how relatively small differences in temperature during the extension phase of diapause can trigger such dramatic changes in genes and pathways, which do not occur during the initiation phase.

Another main finding that arises from our results is that gene expression is not halted at low temperature. Rather, the central gene and signals that participate in embryonic pluripotency are expressed in embryos in diapause at 12 °C, similar to embryos that are not in diapause, but they are downregulated in diapause at 18 °C. Incredibly, other eukaryotic organisms, such as fungi, also show a unique survival mechanism at low temperature by upregulating genes involved in ribosome and energy metabolism, which cannot survive at higher temperature due to upregulation of genes involved in unfolded protein binding, protein processing in the endoplasmic reticulum, proteasome, spliceosome and mRNA surveillance (Su et al. 2016). Taken together, organisms have evolved a cold-adaptation mechanism by not shutting down their molecular machineries at low temperatures.

Finally, our results demonstrate that treating chick embryos in ovo with an exogenous protein source provides a useful means to specifically regulate signaling pathways during embryogenesis (Gallego-Díaz, Schoenwolf, and Alvarez 2002; Levin et al. 1995). Blastulation-stage embryos lost their pluripotency state during the diapause extension phase; however, activation of the BMP4 pathway in these embryos restored the lost pluripotent state, thus highlighting the plasticity of blastodermal cells to regain pluripotency. The induced pluripotency in vivo in blastulating embryos in diapause represents a molecular event that is rarely attainable in vivo in humans, but is widely studied in vitro in mammalian cells (Gallego-Díaz, Schoenwolf, and Alvarez 2002; Tapia et al. 2012) this phenomenon is key to development and regeneration in non-mammalian vertebrates, including teleost fish, urodele amphibians, and lizards (reviewed in Zahumenska et al. 2020).

In summary, this study shows that diapause can be divided into phases which respond differently to the surrounding temperature and affect the embryo’s ability to successfully resume development when returned to incubation. Furthermore, this study identifies a pathway and genes that respond to different diapause phases and temperatures. Verification of these findings revealed the central role of BMP4 in enabling eggs in diapause to successfully resume development by regulating the expression of pluripotency-related genes during diapause. Taken together, our data contribute information on the molecular mechanism underlying the survival of avian embryos during and after diapause, which is likely to provide an evolutionary advantage for avian embryo survival during dormancy in unpredictable environments and for their ability to resume normal development and hatch, accordingly.

## Materials and methods

### Egg collection and blastoderm isolation

Freshly laid broiler eggs (Ross 308) were collected from a local commercial breeder farm. Eggs were stored at 18 °C or 12 °C for up to 28 days with or without treatments as per the experimental design, described previously (Pokhrel et al. 2017). Isolated blastoderms were either placed in RNA Save solution (Biological Industries, Catalog No. 01-891-1A) for RNA stabilization or fixed in 4% paraformaldehyde (PFA) in PBS (Hylabs, Catalog No. BP507/1LD) for further analysis.

### Cell culture

Control and Noggin-secreting CHO cells (Lamb et al. 1993; Sela-Donenfeld and Kalcheim 2002) were cultured in 10-cm plates to 100% cell confluence in 10 mL Dulbecco’s Modified Eagle Media (DMEM; Biological Industries, Catalog No. 010521A) consisting of 10% (v/v) fetal bovine serum (Biological Industries, Catalog No. 040071A), 1% (v/v) L-Glut (GlutaMAX supplement) (Gibco, Catalog No. 35050038) and 1% (v/v) penicillin-streptomycin solution (Biological Industries, Catalog No. 030311B). After cells reached 100% confluence, they were grown in 5 mL of medium for 24 h to obtain a higher concentration of Noggin protein in the conditioned media. The obtained media from control or Noggin-secreting cells were collected and kept at 4 °C as per the experimental design.

### Application of CHO-conditioned media or recombinant BMP4 protein on blastoderms using pluronic gel

CHO-control- and CHO-Noggin-conditioned media were collected and mixed to prepare 15% (w/v) pluronic gel. Pluronic gel suspension was prepared by dissolving Pluronic F-127 powder (Sigma, Catalog No. P2443-250G) in 10 mL protein media extract for 24 h at 4 °C, and protein media extract was then further added to obtain 15% pluronic gel. The gel suspension was prepared at 4 °C because at this temperature, the gel remains in liquid form following dissolution with protein media extract. At temperatures above 10 °C, the gel mixture solidifies. BMP4 recombinant protein (R&D Systems, Catalog No. 314-BP-050) was diluted in 15% pluronic gel (prepared in PBS) to a final concentration of 200 ng/mL, as previously described (Burstyn-Cohen et al. 2004). Pluronic gel suspension was applied on blastulation-stage embryos through a small eggshell window created at the blunt end of the egg through the air sac. The eggshell window was sealed with 2.5-cm wide Leukotape (BSN Medical GmbH, Catalog No. 72668-01). Fresh embryos with CHO-Noggin were additionally stored for 7 days at 12 °C and BMP4-treated embryos were stored for 1 to 3 days at 18 °C according to the experiment. Following treatment, embryos were isolated and placed in RNA Save solution or fixed in 4% PFA for further analysis.

### Application of BMP4-soaked beads on blastoderms

Heparin acrylic beads (Sigma, Catalog No. H5263) were soaked at 4 °C (on ice) for 2 h with BMP4 recombinant protein at a concentration of 1 μg/mL (Trousse, Esteve, and Bovolenta 2001; Weisinger, Wilkinson, and Sela-Donenfeld 2008). Beads soaked in PBS were used as controls. Beads were applied on 27d/18°C embryos through a small eggshell window at the blunt end of the egg through the air sac. Specifically, the beads were implanted in the ventral region of the embryo through a small rupture made by a sharpened needle. The eggshell window was sealed and eggs were held at 18 °C for a further 24 h. Following this treatment, embryos were isolated, fixed in 4% PFA and subjected to in-situ hybridization procedure.

### Real-Time PCR analysis

RNA was extracted from isolated embryos using TRI Reagent (Molecular Research Center, Catalog No. TR-118) and 1 μg of RNA was taken to prepare the cDNA library using the Promega cDNA Synthesis Kit (Catalog No. A5001). Gene expression was quantified by real-time PCR (StepOnePlus Real-Time PCR System, Applied Biosystems) using 1 μL of cDNA, 3.8 μL of molecular biology-grade water (Biological Industries, Catalog No. 018691A), 0.1 μL of each forward and reverse primer, and 5 μL of Applied Biosystems Fast SYBR Green PCR Master Mix (Thermo Fisher Scientific, Catalog No. 4309155). Real-time PCR amplification was performed using the following program: 95 °C for 20 s and 40 cycles of 95 °C for 1 s, then 60 °C for 20 s. Primers used for gene-expression analysis were: GAPDH – Fwd ACCTGCATCTGCCCATTTGA, Rev ACTGTCAAGGCTGAGAACGG; BMP4 – Fwd TCATCCCCAGTTACATGCTG, Rev GCTCTCCAGGTGCTCTTCAT; Id2 – Fwd CTGACCACGCTCAACACAG, Rev TGCTGTCACTCGCCATTAGT; Nanog – Fwd CTCTGGGGCTCACCTACAAG, Rev AGCCCTGGTGAAATGTAGGG; Pax6 – Fwd CCGCACATGCAGACACACATGAAT, Rev TCACTGCCAGGAACTTGAACTGGA.

### Whole-Mount in-Situ Hybridization

Fixed blastoderms were washed with PBT (PBS + 0.01% v/v Tween20 [CAS 9005-64-5, Amresco]), treated with a 0.5% (w/v) solution of deoxycholate (CAS 302954, Sigma) for 8 min, and WMISH was performed as previously described (Weisinger, Wilkinson, and Sela-Donenfeld 2008) with the following digoxigenin (DIG)-labeled riboprobes: Nanog (Lavial et al. 2007), and Id2 (Lorda-Diez et al. 2009). DIG-labeled RNA probes were produced using DIG-UTP by in vitro transcription with T3 RNA polymerase (Promega, Catalog No. P208C) according to the Roche Applied Science’s protocol (Roche Applied Science, Germany). Labeled RNA probes were used for RNA hybridization reaction in whole-mount embryos at an estimated concentration of 1 μg.

### RNA-Seq analysis

RNA-Seq gene-expression profiling of embryos under four different storage conditions (0 day – unstored, 7d/18°C, 28d/18°C and 28d/12°C) was performed. Each group included four biological replicates. Embryos were isolated as previously described (Pokhrel et al. 2017) to obtain 16 individual blastoderms. RNA was extracted from cells with the RNeasy Micro Kit (Qiagen, Catalog No. 74004) using the Qiacube automated system (Qiagen). Quality control for total RNA was performed using TapeStation (Agilent). The RINe value of all samples was in the range of 6.8 to 9.7, with most of the samples presenting RINe values above 8.0. RNA-Seq libraries (NEBNext Ultra II Directional RNA Library Prep Kit for Illumina, New England BioLabs, Catalog No. E7760) were produced from 16 samples according to the manufacturer’s protocol using 200 ng total RNA; mRNA pull-up was performed using the Magnetic Isolation Module (New England BioLabs, Catalog no. E7490). All 16 libraries were mixed at equal molarity in a single tube. The RNA-Seq was generated on an Illumina NextSeq500 system (Catalog No. FC-404-2005), 75 cycles, high-output mode. Quality of single-end reads was assessed using Fastqc (v0.11.5). Reads were then aligned to the Chicken reference genome and annotation file (Gallus_gallus-5.0 and Gallus_gallus.Gallus_gallus-5.0.93.gtf downloaded from ENSEMBL) using STAR aligner (STAR_2.6.0a). The number of reads per gene was counted using Htseq (0.9.1). The descriptive statistical analysis was performed using the DESeq2 R package (version 1.18.1) (Love, Huber, and Anders 2014) and the lists of DEGs were generated. Following data generation, the DEG lists were analyzed using WebGestalt (Liao et al. 2019). Over-representation (enrichment) analysis was used for pathway analysis with the KEGG functional database (Kanehisa 2019; Yi et al. 1999).

### Immunofluorescence

PBS-or BMP4-soaked beads were implanted in the embryos’ trunk neural tube for 24 h. The embryos were then fixed in 4% PFA, rinsed in PBS, permeabilized in PBS + 0.2% (v/v) Triton X-100 (CAS 9002-93-1, Sigma), blocked in 10% normal goat serum in PBS for 2 h and incubated with HNK-1 primary antibody (diluted 1:500; Developmental Studies Hybridoma Bank, AB-531908) overnight at 4 °C. Embryos were washed several times in PBS, and incubated with fluorescent secondary antibody (diluted 1:300 in PBS; goat anti-mouse, Alexa488, Thermo Fisher Scientific, Catalog No. A32723) overnight at 4 °C. Signal was detected in the embryos by fluorescence microscope (model SZX2-ILLK, Olympus; Figure 4-figure supplement 3).

### Western blot analysis

Expression of Noggin in CHO-Noggin cells was examined by extracting total protein from 100% confluent stable Noggin-expressing CHO cells and running it on a western blot. First, the adherent growing cells were washed with 2 mL PBS; 2 mL TrypLE trypsin (Gibco, Catalog No. 12563011) was then added to the plate and incubated for 3 min at 37 °C. The plate was shaken at 1-min intervals to detach the cells. The reaction was stopped with the addition of 3 mL DMEM, and cells were collected in a 15-mL tube (Cell Star, Catalog No. 188261) and centrifuged (model LMC-3000, BioSan) for 5 min at 500*g*. The supernatant was discarded, and the pellet was resuspended in 2 mL PBS, centrifuged for 5 min, and the supernatant was discarded. A 1-mL aliquot of radioimmunoprecipitation assay (RIPA) buffer was added to the cell pellet for 30 min on ice with vortexing (Model FV-2400, BioSan) at 5-min intervals. The cell lysate was centrifuged (Model 5427R, Eppendorf) for 30 min at 4 °C, 14,000*g* and the sample buffer was added to the obtained protein extract. After boiling the mixture at 95 °C for 5 min, the samples were run on the gel. Bands were transferred to a nitrocellulose membrane and treated with Ponceau stain for 1 min to observe the total protein on the membrane. Ponceau was washed from the membrane with DDW and 20 mL of blocking solution was added for 1 h at room temperature (RT). Subsequently, the membrane was incubated overnight at 4 °C with anti-Noggin antibody (diluted 1:2000; Abcam, ab16054). Following three 10-min washes with Tris-buffered saline (1XTBS) consisting Tween (1:1000), 3.5 μL secondary antibody (diluted 1:3000; goat polyclonal anti-rabbit IgG, preadsorbed (horseradish peroxidase [HRP]), Sigma (Merck), SAB3700885) was added in 10 mL of blocking solution and incubated at RT for 1 h with shaking. HRP-conjugated anti-tubulin antibody (diluted 1:1000; Abcam, ab21058) was incubated for 1 h at RT and the signal was detected using SuperSignal West Pico PLUS chemiluminescent substrate (Thermo Fisher Scientific, Catalog No. 34577; Figure 6-figure supplement 5).

### Statistical analysis

Data were analyzed by t-test and one-way ANOVA. Error bars were expressed as mean + SEM. Statistical tests were performed using JMP software (189-2007, SAS Institute Inc.) at a significance level of *P* < 0.05.

---

**Supplemental Figure 1:**
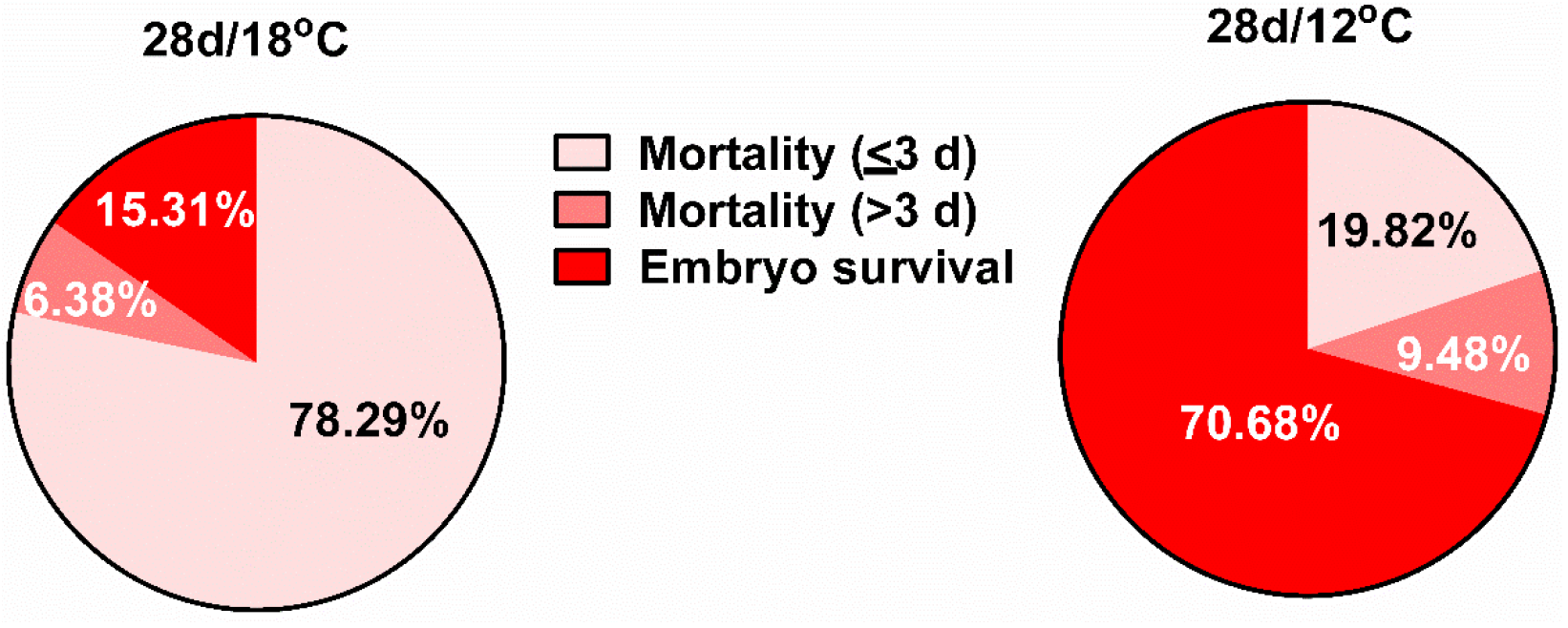
Pie chart showing better survival of embryos (successful resumption of development) following diapause of up to 28 days at 12 °C, compared to 18 °C. Following diapause of up 28 days at 18 °C, most embryos failed to successfully resume development within 3 days of embryogenesis, resulting in 16% embryo survival during hatch. Following 28 days of diapause at 12 °C, early embryo mortality decreased and 71% of embryos successfully resumed development and hatched after 21 days of incubation.

**Supplemental Figure 2:**
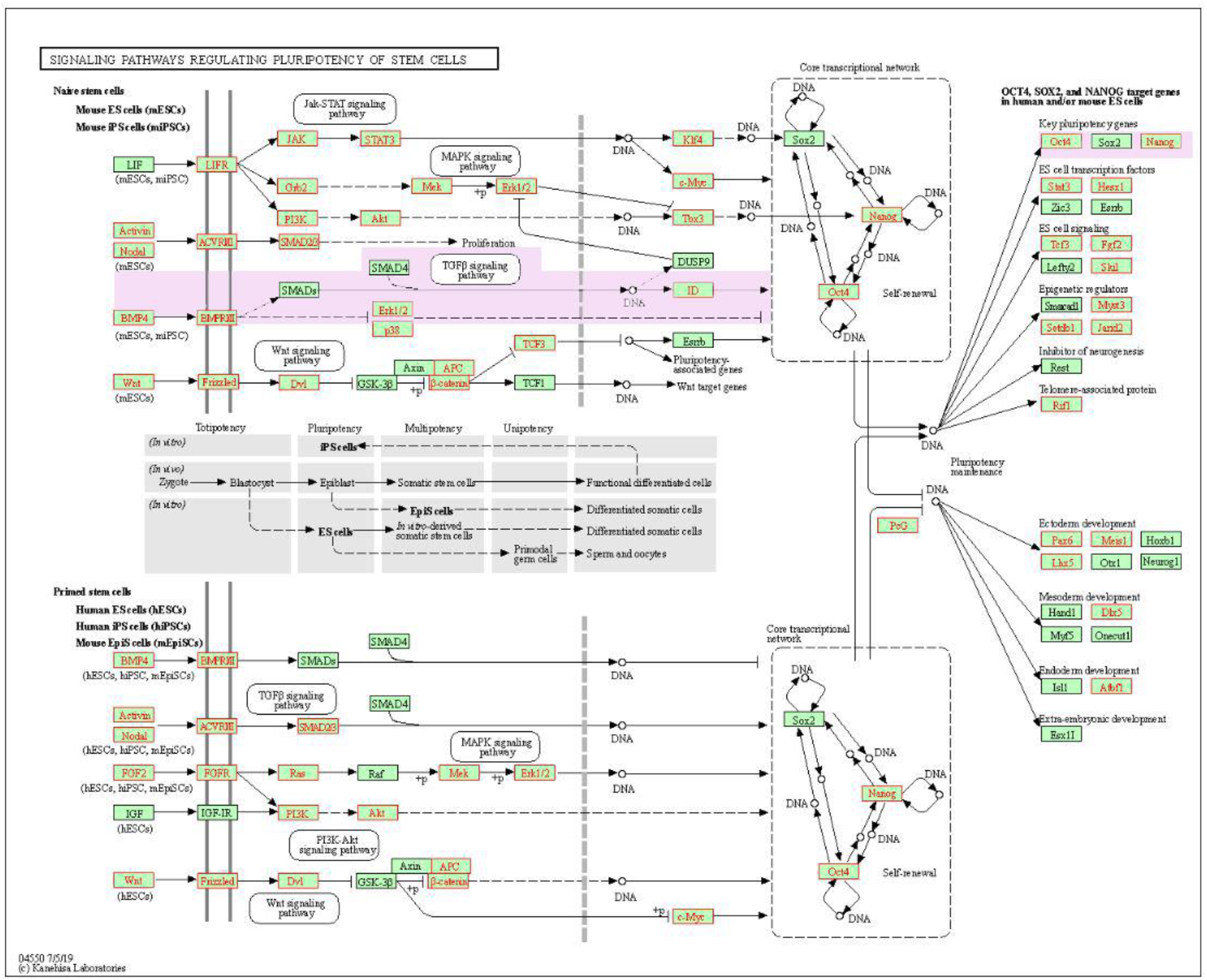
KEGG pathway-enrichment analysis of differentially expressed genes (DEGs) between 28d/18°C and 28d/12°C groups. Analyzing the gene-expression profiles of 28d/18°C and 28d/12°C groups showed enrichment of signaling pathways regulating pluripotency of stem cells in the latter. The enriched gene sets of the TGF-β signaling pathway at 28d/12°C compared to 28d/18 °C are highlighted (light purple) and marked in red, including *BMP4, BMPRI/II, Id* genes, *Oct4*, and *Nanog*. Notably, *BMP4* is upstream of the *Id* genes regulating the core transcriptional network involved in pluripotency, including *Nanog*.

**Supplemental Figure 3:**
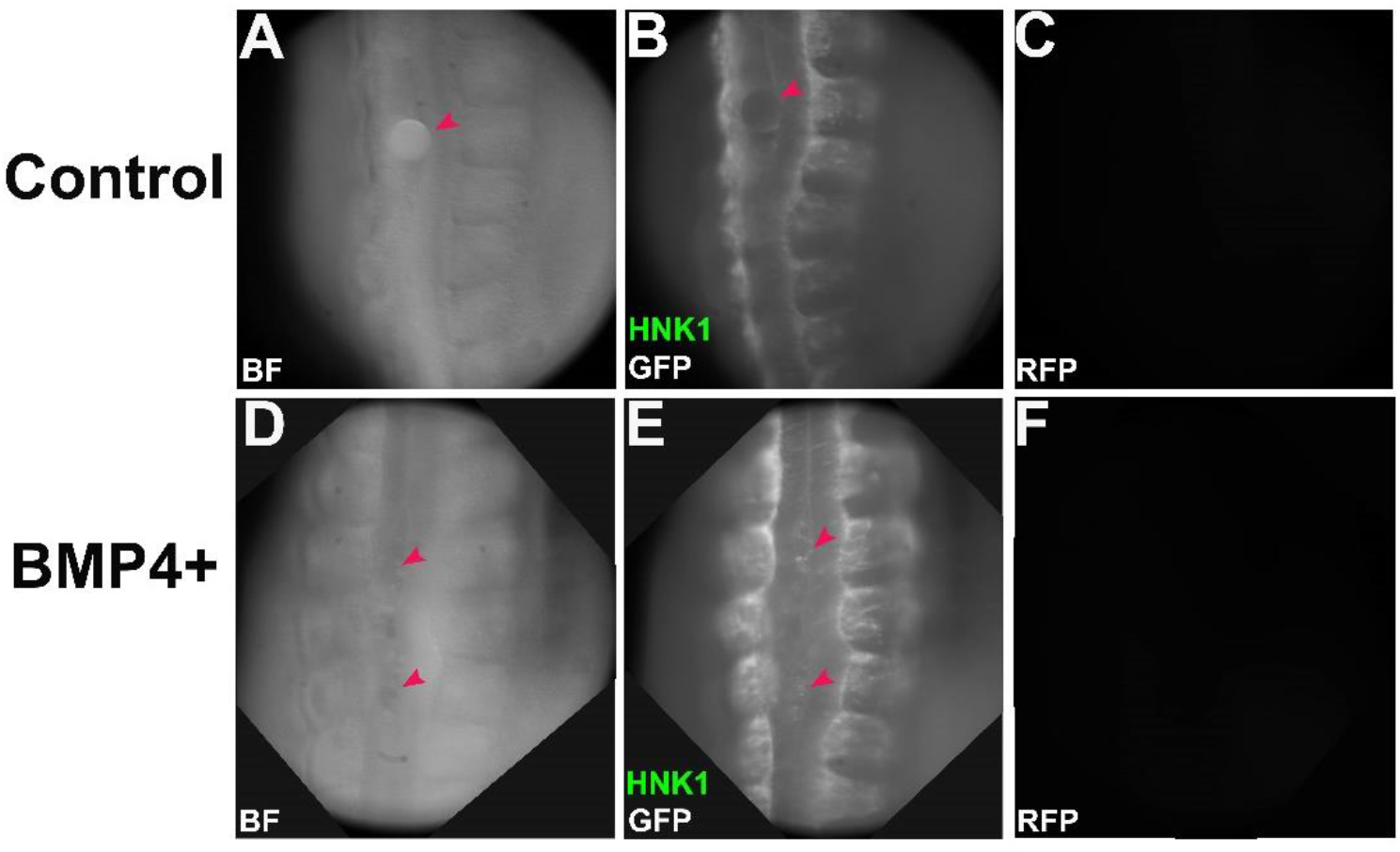
Positive control of BMP4 treatment. Embryos were incubated for 48 h and subsequently treated with beads soaked in either PBS (**A–C**) or recombinant BMP4 protein (**D–F**) for 24 h. Embryos were then fixed in 4% PFA for 24 h and subjected to immunohistochemistry procedure against HNK-1 antibody. (**A–C**) Embryos treated with beads soaked in PBS show HNK-1 staining in the dorsolateral region of the neural tube (indicating neural crest cell delamination) while the mid-dorsal neural tube region does not contain HNK-1-positive cells (**B**, red arrowhead). (**D–F**) Embryos treated with beads soaked with recombinant BMP4 protein show HNK-1-positive cells in both the mid and lateral regions of the neural tube (**E**, red arrowhead), indicating that beads treated with BMP4 recombinant protein can induce neural crest cell delamination in that region. Embryos in panels **B** and **E** were further imaged with RFP filter (**C** and **F**) to confirm specific GFP signal.

**Supplemental Figure 4.**
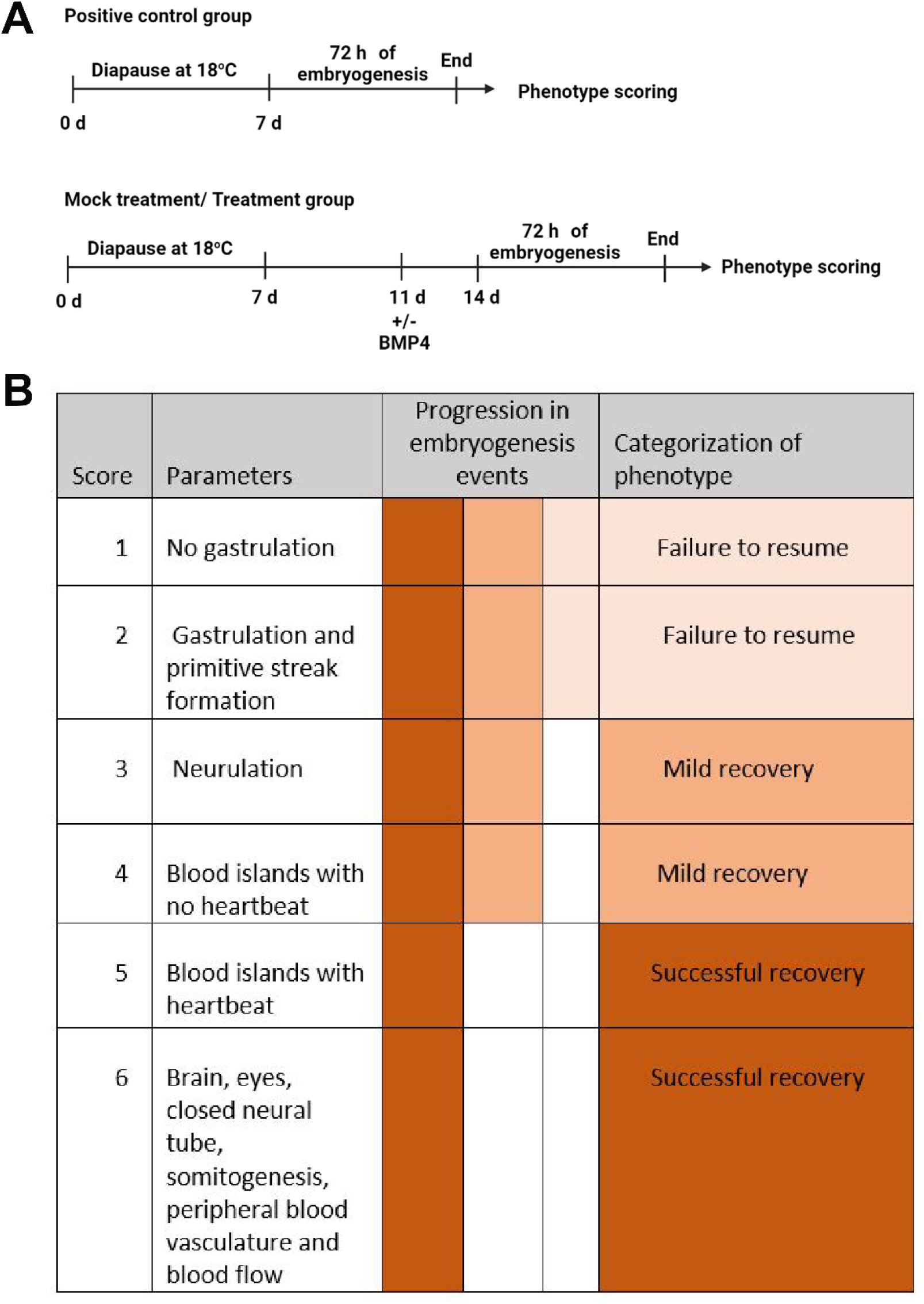
Phenotype scoring of embryos in diapause at 18 °C that successfully resumed development (SRD) after 72 h of incubation. (**A**) Experimental design for phenotype scoring of embryos in diapause with (+BMP4) or without (-BMP4) BMP4 application. Embryos stored for 7 days at 18 °C were used as positive controls, as they showed better SRD. Mock control and treatment groups included embryos that underwent diapause for 11 days at 18 °C and were further stored for 3 days with (+BMP4) or without (-BMP4) BMP4. All three groups: positive control, mock control and treatment group, underwent 72 h of embryogenesis and then their phenotype was scored. (**B**) Following 72 h of embryogenesis, based on morphological features, the phenotypes of positive control, mock control and rescued embryos (treatment group, +BMP4) were categorized as failure to SRD (failure to resume), mild recovery and successful recovery phenotypes. After 33–36 h of normal embryogenesis, scattered blood islands are already formed. Therefore, embryos that did not form blood islands after 72 h were defined as failure to SRD (no SRD characteristics). Embryos that formed blood islands without a heartbeat were defined as mild recovery, and, those with a heartbeat and peripheral blood vasculature, clear blood flow, brain, eyes, closed neural tube and somitogenesis, were defined as successfully recovered. After categorizing the embryos, successive development events were scored from 1 to 6 and then the phenotypes were analyzed.

**Supplemental Figure 5.**
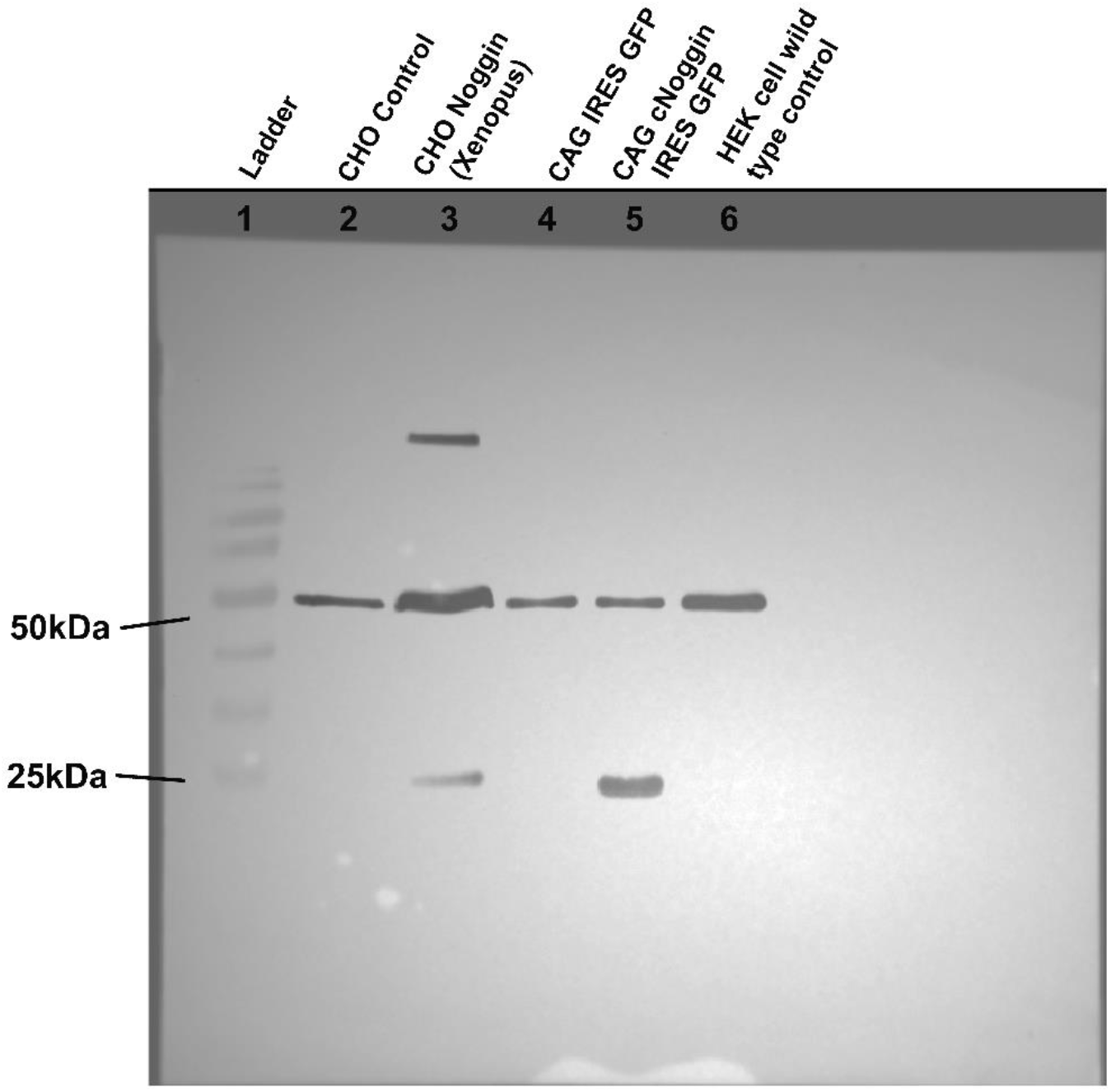
Western blot analysis confirming Noggin protein production by CHO-Noggin cells. Lanes: 1 – ladder, 2 – CHO Control, 3 – CHO Noggin (*Xenopus*), 4 D CAG IRES GFP, 5 – CAG cNoggin IRES GFP (positive control), 6 – HEK wild-type control cell. Expected band size for anti-tubulin and anti-Noggin antibody is 50 and 26 kDa, respectively.

## Notes

**Competing Interest Statement:** The authors declare no conflict of interest.

### Competing Interest Statement

The authors have declared no competing interest.

