## Supplementary figures and images for "How do avian embryos resume development following diapause? A new role for TGF-β in regulating pluripotency-related genes"

### Figure 1-figure supplement 1.tif

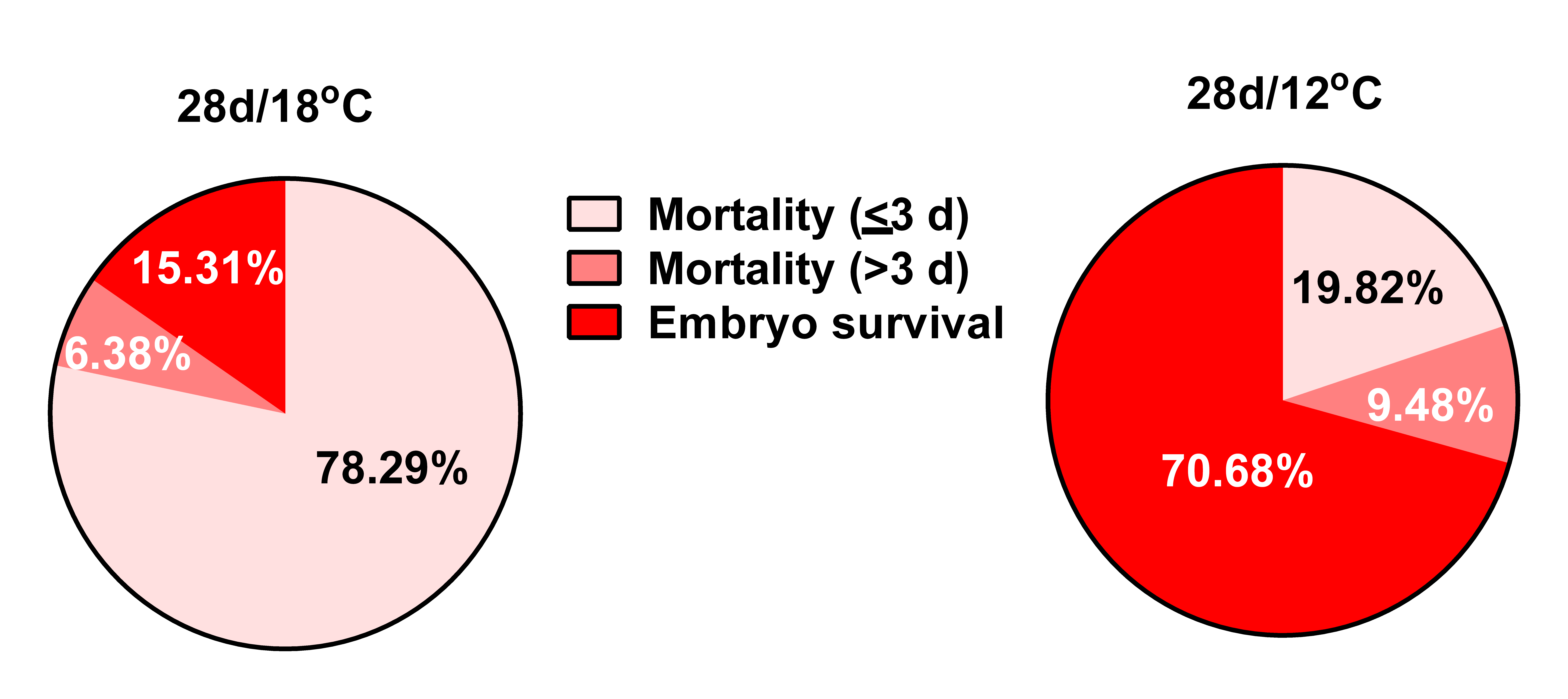

### Figure 2-figure supplement 2.tif

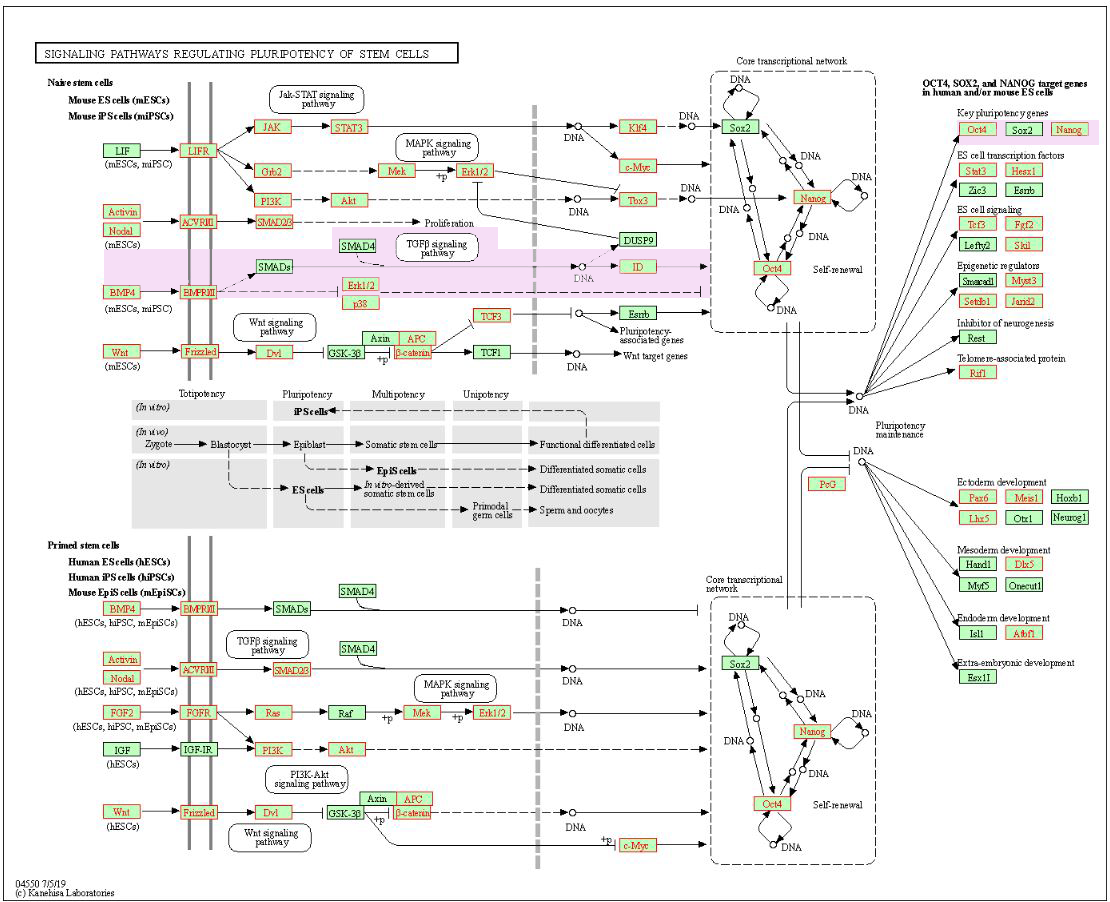

### Figure 4-figure supplement 3.jpg

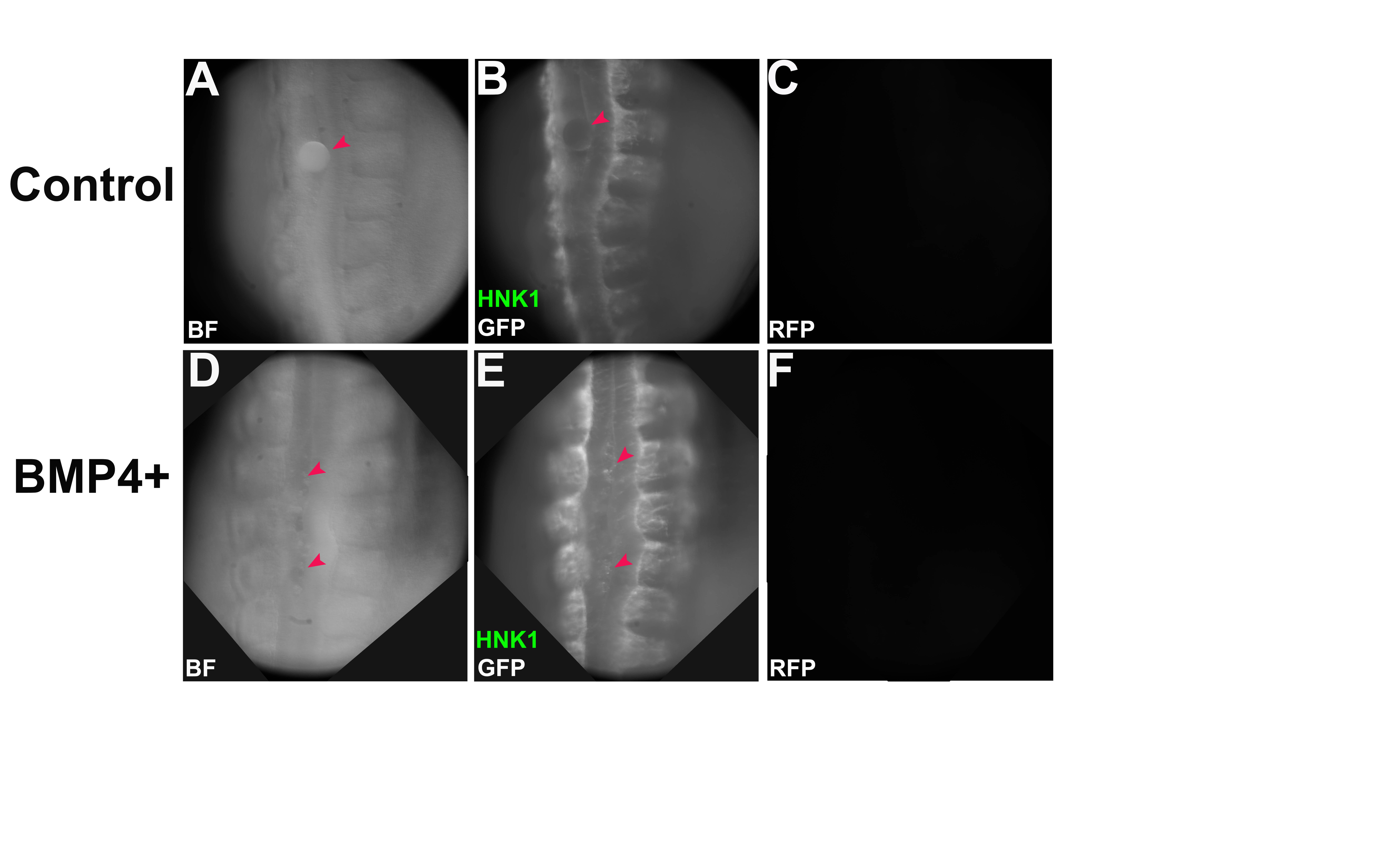

### Figure 6-figure supplement 5.jpg

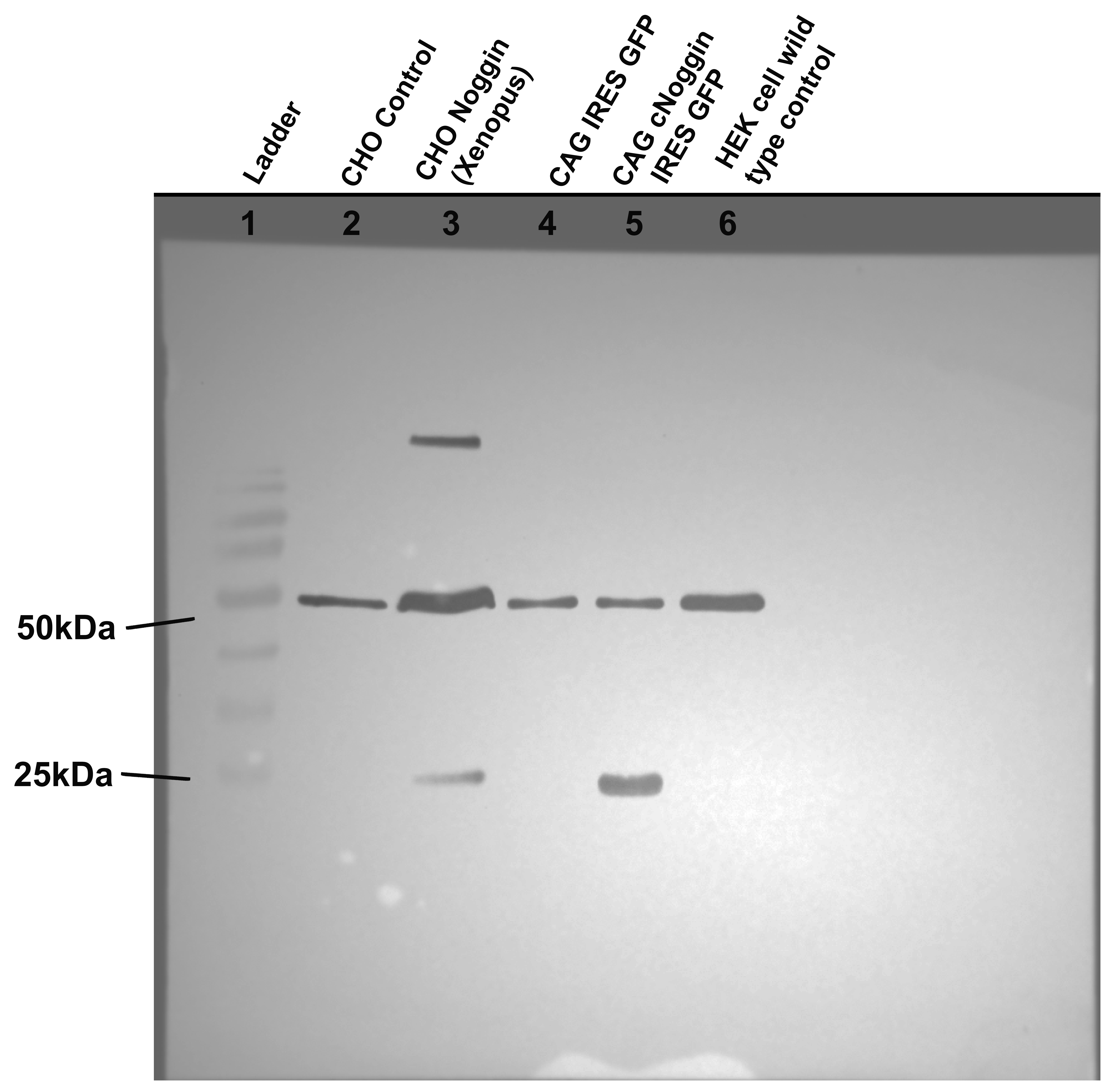

### Figure 8-figure supplement 4.jpg

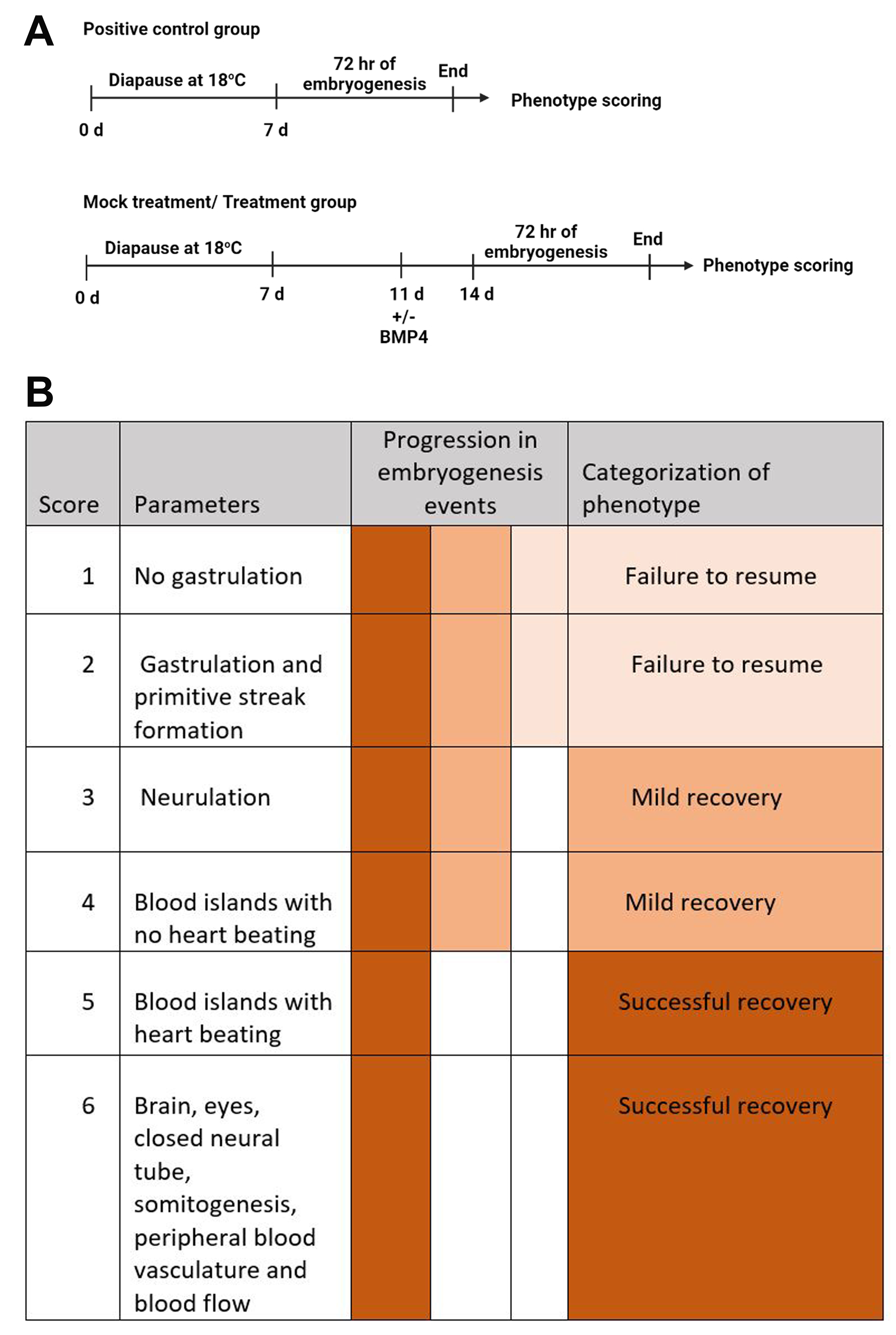
